# Discovering Broader Host Ranges and an IS-bound Prophage Class Through Long-Read Metagenomics

**DOI:** 10.1101/2025.05.09.652943

**Authors:** Jakob Wirbel, Angela S. Hickey, Daniel Chang, Nora J. Enright, Mai Dvorak, Rachael B. Chanin, Danica T. Schmidtke, Ami S. Bhatt

**Affiliations:** Division of Hematology, Department of Medicine, Stanford University, Stanford, CA; Department of Genetics, Stanford University, Stanford, CA; Department of Bioengineering, Stanford University, Stanford, CA; Department of Biology, Stanford University, Stanford, CA; Department of Microbiology and Immunology, Stanford University, Stanford, CA

**Keywords:** Microbiome, Phages, Long-read sequencing, Metagenomics, Insertion sequences

## Abstract

Gut bacteriophages profoundly impact microbial ecology and human health, yet they are greatly understudied. Using deep, long-read bulk metagenomic sequencing, a technique that overcomes fundamental limitations of short-read approaches, we tracked prophage integration dynamics in 12 longitudinal stool samples from six healthy individuals, spanning a two-year timescale. While most prophages remain stably integrated into their host over two years, we discover that ∼5% of phages are dynamically gained or lost from persistent bacterial hosts. Within the same sample, we find evidence of population heterogeneity in which identical bacterial hosts with and without a given integrated prophage coexist simultaneously. Furthermore, we demonstrate that phage induction, when detected, occurs predominantly at low levels (1-3x coverage compared to the host region). Interestingly, we identify multiple instances of integration of the same phage into bacteria of different taxonomic families, challenging the dogma that phage are specific to a host of a given species or strain. Lastly, we describe a new class of phages, which we name “IScream phages”. These phages co-opt bacterial IS30 transposases to mediate their integration, representing a previously unrecognized form of phage domestication of selfish bacterial elements. Taken together, these findings illuminate fundamental aspects of phage-bacterial dynamics in the human gut microbiome and expand our understanding of the evolutionary mechanisms that drive horizontal gene transfer and microbial genome plasticity in this ecosystem.

## Introduction

Bacteriophages are the most abundant biological entity on earth and play crucial roles in shaping microbial communities^1^. Classically, phages are either lytic or exist in an integrated state in their bacterial host (lysogen). The deep study of several model integrated phages (such Lambda and Mu) has built the framework for our understanding of lysogens, and has also informed many aspects of molecular biology^2^. More recently, advances in metagenomic sequencing and improved analysis tools have shown that lysogens are much more diverse, prevalent and abundant in human microbiomes than previously appreciated^3–7^.

Multiple lines of evidence suggest that the majority of phages in the human gut exist as integrated prophages^8^, even though alternative lifestyles, such as phage plasmids or carrier states also exist^9–11^. Typically, prophages insert their genome into the host chromosome using dedicated integrases of three different classes: tyrosine recombinases (YRs), small and large serine recombinases (SRs), and DDE recombinases. Characterization of these integration or recombination mechanisms has formed the basis of many modern molecular tools, such as Cre recombinase, which is used extensively for creating transgenic cells and animals^12^. By integrating into the host chromosome, phages can ensure replication and vertical transfer within their host when conditions are not optimal for lytic replication. Hosts may derive a fitness benefit from prophage-encoded genes such as antibiotic resistance cassettes or virulence factors^13,14^ but run the risk that prophages may re-enter the lytic cycle and thus kill their host. Prophages can accumulate mutations over time, leading to decay of mobilization machinery^15^, whereas other prophage genes may be adopted and maintained by their bacterial host^16^. Similarly, prophages may incorrectly package host genes upon induction, thereby mediating horizontal gene transfer (HGT) between related bacterial strains or species, through processes such as generalized and specialized transduction^17,18^.

Prophages can have profound impacts on their bacterial host and the microbial community at large, yet it remains challenging to study phages and hosts within the same sample. Distinguishing between phage and bacterial regions of the chromosome has not been straightforward, prompting researchers to employ virus-like particle (VLP) sequencing to enrich for viral sequences^19^. Studies of the human gut virome using this method have suggested that phages in the gut are relatively stable, persisting within their human superhost over one year^20,21^. VLP-based studies form the basis of our knowledge of phage abundance and stability in the human gut microbiome, but this technique has two notable limitations: focusing on VLPs (1) overlooks prophages that are not actively replicating and producing phage particles at the time of sampling and (2) there is little direct evidence of which prokaryotic host(s) each phage infects. Bulk metagenomics, where viral and prokaryotic genomes are sequenced simultaneously, somewhat addresses these challenges given novel phage prediction tools^22–24^. However, most bulk studies are performed with short-read sequencing. This approach is limited in its ability to resolve repetitive elements, posing many challenges when studying phages derived from complex communities. For example, phage genomes can have large regions of similarity (phage genomic mosaicism^25^) leading to fragmented assemblies and incomplete genomes. Similarly, if a given prophage is integrated into multiple hosts, short-read assembly would be unable to resolve its hosts, complicating further investigation.

Long-read metagenomic sequencing can address these challenges by yielding more contiguous assemblies, potentially resolving individual phage genomes and the hosts of integrated prophages. Previous studies have demonstrated advantages of long-read sequencing for the study of phages: one study using long-read VLP sequencing was able to resolve more complete viral genomes and identified structural variations within phages^3,26^. Another study reported improved prophage and CRISPR spacer assembly with long-read sequencing, capturing individualized phage populations that were stable over a ten-day period^27^. Improved CRISPR spacer assembly and similarity-based approaches have enabled host prediction from complex communities, suggesting extremely broad host range for some phages^3,4,28^, which challenges the notion of exquisitely narrow host range^29^. However, CRISPR spacers are typically very short (20-50 nucleotides), potentially over-estimating host range, necessitating further molecular evidence^30^. Accurate assembly of prophages into their native host through long-read assembly would provide direct evidence of infection, enabling the study of specific phage-host relationships. With improvements in accuracy^31^ and cost, long-read sequencing is becoming increasingly accessible for longitudinal metagenomics and is poised to improve the detection of prophages and interactions with their hosts.

Here, we sought to examine the relationship between prophages and their hosts in the healthy human gut using long-read sequencing. To do so, we generated a deep, long-read, longitudinal metagenomic dataset from stool samples of six individuals who were studied over a two-year time period. This enabled us to study the dynamics of integrated prophages in multiple individuals across longer timescales than have been previously reported. While we find a small fraction of phages to be gained or lost over two years, most phages seem to be stably integrated into their bacterial host. Prophages appear to induce at predominantly low levels, with some phages and their hosts existing in mixed populations (induced phage, integrated phage, host without phage) at the same time. A small number of phages assemble into multiple and taxonomically diverse host contexts, providing strong evidence for broad host range. Unexpectedly, we also identify a group of related prophages, IScream phages, that do not encode their own canonical integrase, but instead have likely domesticated bacterial insertion sequence (IS) elements for their integration and excision machinery. In this study, we demonstrate how longitudinal long-read metagenomics can help elucidate diverse aspects of prophage biology in human microbiomes.

## Results

### Generating a long-read, longitudinal metagenomic dataset to study integrated phages

To learn more about the biology of integrated phages and their activity over time, we collected stool samples from six healthy adults at two timepoints, two years apart (T1 and T2). We then generated long-read metagenomic DNA sequencing data from these samples on the Oxford Nanopore Technologies (ONT) platform to a depth of ∼30 billion bases (Gb) (see **Fig. 1a, SFig. 1, STable 1**). All samples were also sequenced by short-read shotgun sequencing using the Illumina platform to a depth of 6 Gb. To be able to compare between short- and long-read sequencing without the bias of differences in sequencing depth, we subsampled our long-read data to the mean of the short-read data (see **Methods**). After quality control and host-read removal, short reads were assembled with Megahit^32^ and long reads with metaFlye^33^, followed by binning into metagenome-assembled genomes (MAGs) for both assemblies (see **Methods**). As expected, the long-read assemblies exhibited higher contiguity with a higher mean contig N50 (255.5kb for long-read vs 7.8kb for short-reads) and lower number of contigs within corresponding high-quality bins (p=2.2e-16, paired Wilcoxon test, see **SFig. 1**). In contrast to gene calling challenges due to high error rates from previous ONT chemistries^34,35^, the average length of predicted genes was similar across both types of assemblies (see **SFig. 1**). This demonstrates that ONT reads generated with the latest chemistry (R10.4, median read quality ∼Q20) can result in accurate assemblies without the need for short-read polishing^31^. Taken together, our long-read assemblies exhibited much higher contiguity than the short-read-based assemblies, without sacrificing quality.

**Figure 1:**
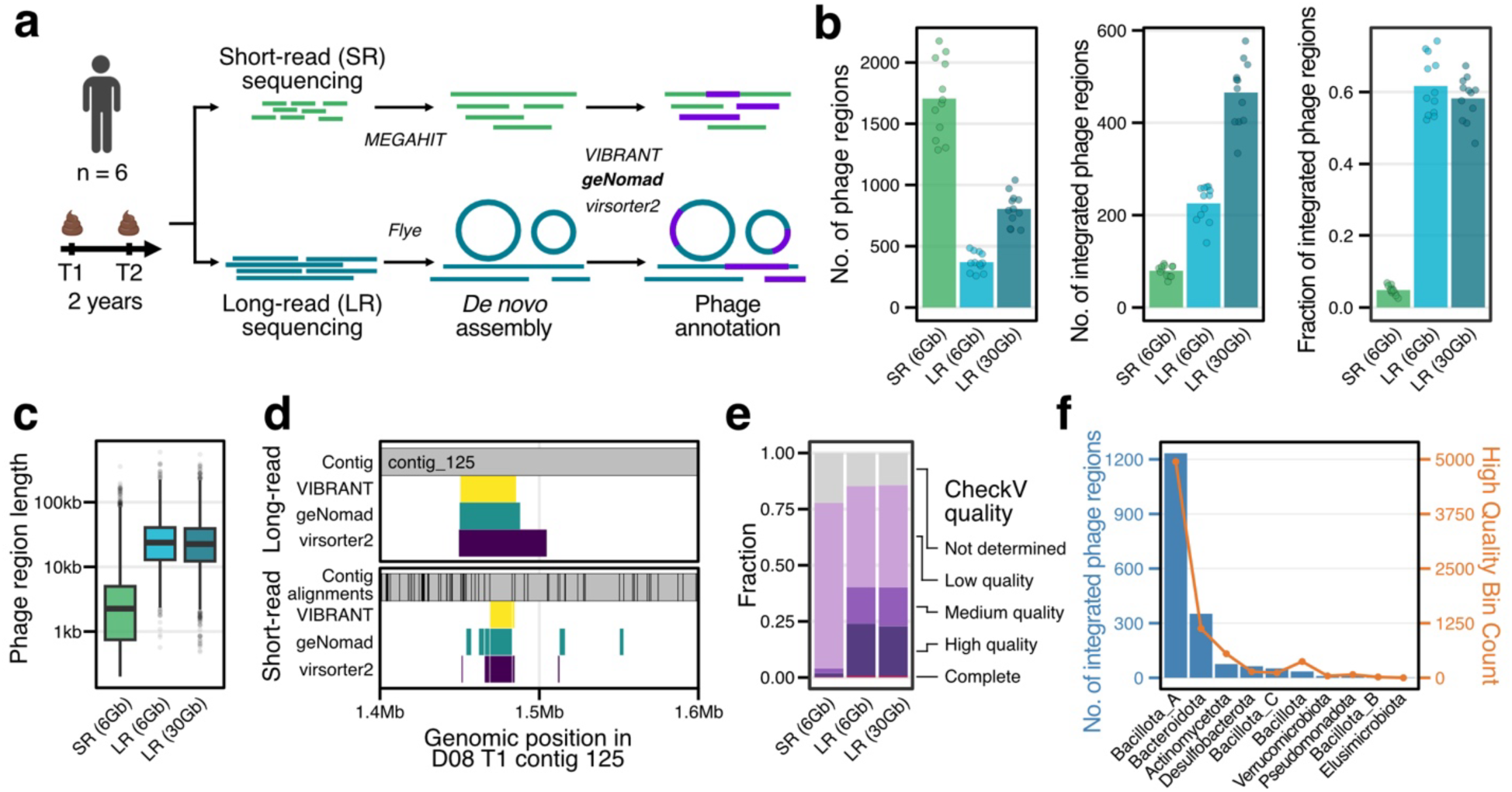
Improved detection of integrated prophages with long-read sequencing. **a)** Schematic of the analysis workflow. DNA from two stool samples collected ∼2 years apart for 6 healthy individuals were sequenced using both short-read (Illumina) and long-read (Oxford Nanopore) platforms. MEGAHIT and metaFlye were used to assemble the short and long reads, respectively. Phages were then predicted from both assemblies using VIBRANT, geNomad, and virsorter2. **b)** Number of phage regions, number of integrated phages, and the fraction of integrated phages predicted by geNomad for the assemblies from short-read, subsampled long-read (6Gb), and deep long-read (30Gb) sequencing. Bars show the mean over all samples and points indicate the values for individual samples. **c)** Length of phage regions is shown for short-read (6Gb), subsampled long-read (6Gb), and deep long-read (30Gb) assemblies. Boxplots show the interquartile ranges (IQRs) as boxes, with the median as a black horizontal line, whiskers extending up to the most extreme points within 1.5-fold IQR, and outliers indicated as dots. **d)** Example for fragmentation with short-read sequencing. Filled boxes indicate sequence regions that are predicted to be phage by the different tools. In the lower panel, grey boxes indicate alignments of short-read contigs against the subsampled long-read assembled contig. **e)** Mean fraction of CheckV quality annotations across samples for phages from the short- and long-read assemblies. **f)** Blue bars show the number of phage regions found integrated in each bacterial phylum, combined across all samples, assessed on the deep (30Gb) long-read assemblies. The number of high quality bins assigned to each bacterial phylum, summed across all samples, is represented by orange points (see right y-axis).

### Improved detection of integrated prophages in long-read assemblies

To compare phages in both short and long-read assemblies, we used the computational phage prediction tool geNomad^24^, which combines alignment-free and gene-based models for viral prediction. Across all samples, geNomad found more phage regions in the short-read compared to the downsampled long-read assemblies. However, the percentage of integrated phages was lower (∼10% in short-read vs ∼60% in long-read assemblies, see **Fig. 1b**). Finding a majority fraction of integrated prophages in the long-read assemblies is consistent with the expectation that most phages in the human gut are thought to be integrated prophages^8^. These results were recapitulated using phage annotations from two other commonly used prediction tools: VIBRANT^22^ and virsorter2^23^ (see **SFig. 2**).

Since the average length of predicted phage regions was much smaller in short-read than in long-read assemblies (see **Fig. 1c**), we hypothesized that a single phage region predicted in the long-read assemblies might be represented by multiple smaller phage regions in the short-read assembly. We therefore mapped the short-read contigs to the long-read assemblies and compared phage annotations across sequencing methods. As an illustration, **Fig. 1d** shows the phage predictions and alignments of short-read contigs against a single long-read contig containing a 50kb region that is predicted by geNomad, VIBRANT and virsorter2 to be an integrated prophage. This region recruits alignments from multiple phage contigs in the short-read assembly, but also from contigs not predicted to be phage, indicating likely mis-annotations (see also **SFig. 3**). Overall, the majority (82%) of long-read integrated phages were found to be fragmented in the corresponding short-read assemblies (see **SFig. 3**, **Methods**). Consistent with this, a higher fraction of long-read phages were classified as medium quality or higher by CheckV^36^, which assesses genome completeness and presence of phage hallmark genes (see **Fig. 1e**). These results demonstrate the superiority of long-read metagenomics for the accurate and complete assembly of integrated phages.

In the following sections, we will focus on the deep (30Gb) versus the downsampled (6Gb) long-read sequencing assemblies as we detected more phage regions with the same quality at this depth (see **Fig. 1b,c,e**).

Since long-read sequencing allowed us to assemble the majority of phages as integrated elements into their bacterial host, we could directly determine the taxonomy of the bacterial hosts of each of the integrated phages from our assemblies. To do so, we used taxonomic assignments of the genes in the surrounding host region to find a consensus host taxon (see **Methods**). We found a high concordance of host assignment between this approach and existing host prediction methods^37^ (∼90% agreement up to the family level, see **SFig. 4**). To ensure accurate host identification, we moved forward with a set of high confidence host assignments with consensus of high-quality bins and gene-level taxonomic assignment. The majority of integrated phages were found in the phyla *Bacillota A* and *Bacteroidota*, which reflects the number of high-quality bins present across samples (see **Fig. 1f**). In summary, long-read metagenomic sequencing and assembly improved the detection of integrated prophages and their hosts.

### Integrated prophages are mostly integrated in a single locus and have low rates of induction

As we had samples from the same individuals that were two years apart, we sought to determine whether phages were gained or lost independent of their bacterial host over time. In this analysis, we also sought to determine whether phages could be found in different genetic loci within the same host or in different hosts, altogether. To compare identical phages between the two timepoints for each individual, we created a non-redundant set of phages from a given sample. Specifically, we clustered all integrated phages within the same individual at 99% identity and 90% genome coverage, while taking their surrounding host regions into account (see **SFig. 5** and **Methods**). We then used this information to quantify the number and types of phages that were found in different genomic loci.

Overall, 35% of phages were found in the same position within their bacterial host (host context) at both timepoints. 56% of phages, along with their hosts, were assembled in a single timepoint only, potentially due to changes in the overall microbial community in the gut (**Fig. 2a**). Dynamic phage clusters, meaning phages that were lost or gained while the host remained persistent, were rather rare (∼5% of clusters). Examples of phages being lost or gained between T1 and T2 in an otherwise stable bacterial host are provided as sequencing coverage plots (see **Fig. 2b & 2c**).

**Figure 2:**
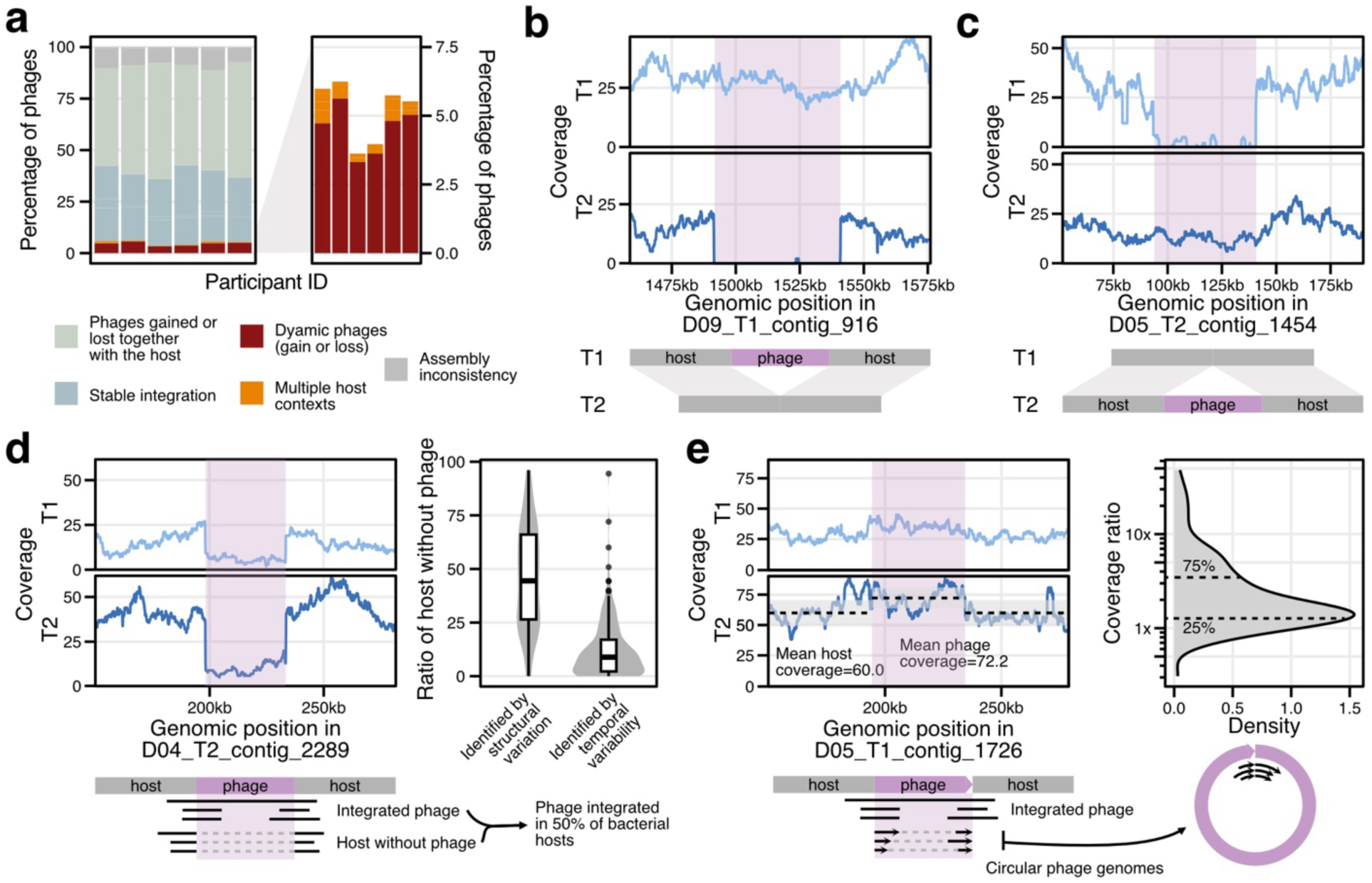
Phage dynamics and population heterogeneity in the gut. **a)** Percentage of high-identity phage clusters and their classification (see **SFig. 5**) per individual. The right panel zooms in on the fraction of clusters that are dynamic or found in multiple host contexts. **b)** Coverage plot supporting the loss of an integrated phage in T2 for individual D09. **c)** Coverage plot supporting the gain of an integrated phage in T2 into a host that is already present in T1. **d)** Coverage plot showing population heterogeneity in terms of integration for a prophage in individual D04. Based on reads supporting the integrated phage or host without phage (see schematic below), the ratio of hosts without phage was calculated. In the right panel, this ratio is shown as a boxplot for all phages with structural variation evidence in the same timepoint and for all phages with structural variation evidence in the other timepoint (identified by temporal variability). Boxplots show the interquartile ranges (IQRs) as boxes, with the median as a black horizontal line, whiskers extending up to the most extreme points within 1.5-fold IQR, and outliers indicated as dots. **e)** Coverage plot for a phage with structural variation evidence for circular phage genomes (see schematic below). The mean coverage of the phage and host region are indicated by dashed black lines (the grey area shows the mean plus/minus one standard deviation). In the right panel, the coverage ratio between phage and surrounding host region is shown as a density plot with the 25% and 75% percentile indicated by dashed black lines.

Interestingly, we also observed drops of coverage for phages that were assembled as stably integrated, suggesting the presence of both hosts with an integrated phage (lysogens) and hosts without an integrated phage (naive hosts) at the same time. To more accurately quantify these cases, we used Sniffles2 to identify structural variations (SVs) from reads mapped across timepoints, which allowed us to find read support for deletions (naive hosts) when the lysogen was assembled (see **Methods**). This approach also revealed the boundaries of the phage genome with base-pair precision, as we were able to discern the exact integration sites (see **SFig. 6**). In total, we found that lysogens and naive hosts coexisted in ∼7% of the cases. We then quantified the proportion of lysogen to naive hosts, and observed a wide distribution of values for this ratio, indicating that prophage prevalence within a population can be highly heterogeneous (see **Fig. 2d**). In a subset of cases (∼40%), we could detect the deletion SV (which corresponds to the naive host) only through reads from the other timepoint, meaning that the proportion of naive hosts within the population changed over time. Using the exact boundaries detected this way, we often observed a small fraction of reads supporting the existence of naive hosts in the original timepoint, despite falling below Sniffles2 detection thresholds (see **Fig. 2d**). This indicates that heterogenous prophage prevalence, where lysogens and naive bacterial hosts can co-exist, might be more common than we are able to detect.

In addition to the detection of naive hosts, SV calling also resulted in the detection of circular phage genomes. Circularization of phage genomes can occur during prophage induction; thus, this allowed us to measure the induction rate of some of the integrated phages *in situ*. In most cases, the presence of circular phage genomes was not concomitant with an appreciable increase in genome coverage compared to the surrounding host region (see **Fig. 2e**). Instead, we observe the majority of induced phages to be present at 1x to 3x coverage compared to the surrounding host region. This is consistent with low level phage induction as opposed to large lytic replication bursts (>10x), the latter of which seems to be the exception in the gut^38^. Curiously, some phages with evidence of circular genomes exhibited coverage ratios lower than 1x compared to the host region, which could be explained by the simultaneous presence of lysogens, naive hosts, and induced circular phage genomes (see **SFig. 6**).

Overall, our data suggest that integrated phages are relatively stable in their hosts over a 2-year timeframe, with few cases of novel integration of a phage or phage loss. Additionally, both phage integration and phage induction are subject to substantial population-level heterogeneity in the gut, with low average rates of phage induction.

### Evidence for broad prophage host range

In addition to detecting phage loss and gain, we are able to find the same prophage in multiple host contexts within and across timepoints (see **Fig. 2a**). In one example, we observe the same prophage (99% average nucleotide identity (ANI)) found stably integrated into two different host contexts of the same host (*Alistipes putredinis*), supported by strong coverage across the region at both T1 and T2 (see **Fig. 3a**). Closer investigation of the two *A. putredinis* bins across timepoints revealed high similarity (98.71% ANI), yet extensive structural differences including deletions, insertions, and recombination (see **SFig. 7**). These observations suggest that a new strain of the same bacterial species colonized this individual during the sample period. This new strain was infected with (or already carried) the same phage, which integrated into a different genomic location (see **SFig. 7**). In another case, we again found the same phage stably integrated in two host contexts within the same individual (see **Fig. 3b**); the hosts are of the same taxonomic family (*Lachnospiraceae*) but differ at the genus level (*Ventrimonas* and *Clostridium Q*). While the host range of most phages is believed to be restricted to single species or even strains, previous computational predictions based on CRISPR spacer analysis had suggested that some phages might have broader host range^3,4,28^. Here, we provide strong, direct evidence for broad host range via the observation of integration of identical phages into taxonomically distant hosts.

**Figure 3:**
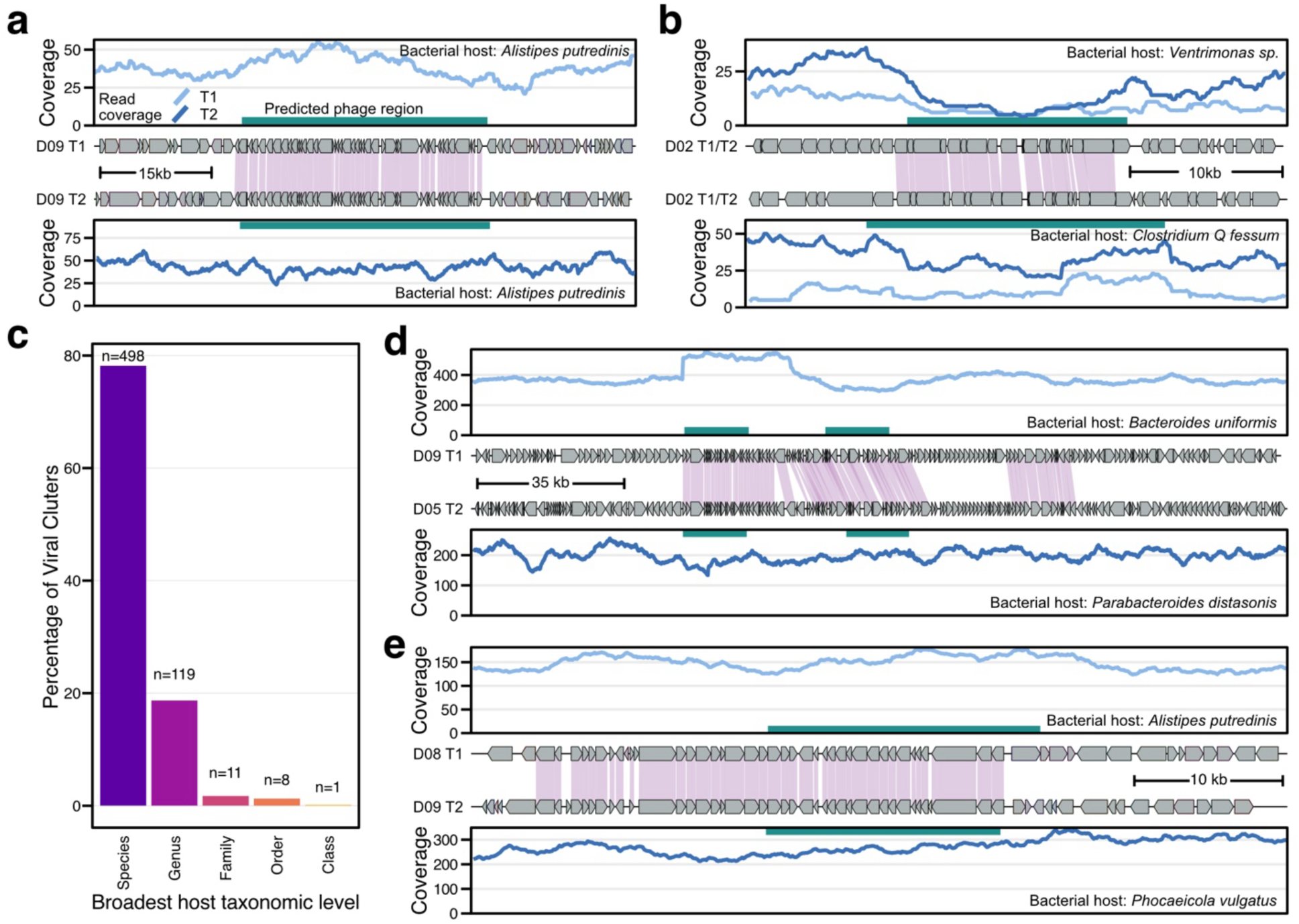
Long-read assemblies provide evidence for a broad host range for integrated phages. **a)** Coverage and synteny plot supporting the shared phage region between two *A. putredinis* contigs assembled in the same individual (D09) in T1 and T2, respectively. For all plots of this type, the phage region predicted by geNomad is indicated by a green bar. **b)** Coverage and synteny plot showing a shared phage region between a *Ventrimonas sp.* contig and a *Clostridium Q fessum* contig, both assembled in T1 and T2 in individual D02. **c)** Percentage of all viral clusters with members that are restricted to the species, genus, family, order, or class taxonomic level. The number of clusters present in groups above species-level is noted above each bar. **d)** Coverage and synteny plots supporting shared genomic regions between a *B. uniformis* contig assembled in T1 from individual D09 and a *P. distasonis* contig assembled in T2 from individual D05. **e)** Coverage and synteny plots showing a shared phage region between a *A. putredinis* contig assembled in T1 from individual D08 and a *P. vulgatus* contig assembled in T2 from individual D09. The shaded regions in panel **a, b, d,** and **e** indicate amino acid similarity greater than 80%.

Since only a few phages within the same individual were present in multiple host contexts (N=10 on average), we next assessed whether closely related phages were found in different hosts across individuals. We used standard clustering cutoffs^36^ for viral species of 95% identity and 85% coverage to cluster all phages across all individuals. Using our high confidence host identification approach, we determined the broadest host taxonomic level that was shared within a phage cluster. As expected, a majority of phages were restricted to the species level (78%), with just under 20% being restricted to the genus level (see **Fig. 3c**). We did find evidence for a small number of phages that demonstrate broad host range: 11 phage species were restricted to the family level and eight were restricted to the order level (see **STable 2**). A single example of a phage species restricted to the class level was insufficiently supported by read coverage (see **SFig. 8**). Six out of eight phage species restricted at the order level were found in the *Bacteriodales* order. In one example, a highly similar phage region (∼50kb) was found integrated in *Bacteroides uniformis* (Family: *Bacteroidaceae*) in one individual and in *Parabacteroides distasonis* (Family: *Tannerellaceae*) in another, well supported by high coverage (see **Fig. 3d**). This phage cluster had a total of seven members from four individuals with hosts found in multiple species within these two families (see **SFig. 8**), revealing integration into variable host contexts. In the two assemblies shown in **Fig. 3d**, we also observed a smaller element shared in both assemblies, possibly representing another mobile element shared across bacterial families. As a last example, we found a phage region shared across two individuals between the *Bacteroidaceae* (Species: *Phocaeicola vulgatus*) and *Rikenellaceae* family (Species: *Alistipes putredinis*, see **Fig. 3e**). The geNomad prediction does not encompass the complete shared region, suggesting that the actual phage region is larger than predicted.

Taken together, these examples provide strong, assembly-level evidence for a broad host range of some phages, mostly infecting hosts of the *Bacteroidales* order.

### IScream phages

In most cases, prophages carry their own DNA integration enzymes (integrases) to enable mobilization of their genomes. When examining the set of prophages with exact genome boundaries from SV evidence (n = 569), we found 359 phages with tyrosine- and 165 phages encoding serine-integrases. 124 had neither tyrosine- nor serine-integrases. As Mu and a small number of other phages use a DDE integrase^39^, we queried the entire set of 569 prophages with exact genome boundaries to find those that contained a DDE integrase. We found 23 prophages that encoded DDE integrases; however, DDE-type enzymes were mostly found in phages that contained integrases of other types as well (n = 20; see **Fig. 4a**). These DDE enzymes resemble those found in bacterial insertion sequences (ISs), which are selfish genetic elements consisting of transposase enzymes flanked by inverted repeats with the ability to mobilize their own sequence^40^. As most of the identified DDE-type enzymes co-occurred in phages with another integrase, these likely represent IS elements that had inserted into the phage genome instead of genuine mobilization machinery.

**Figure 4:**
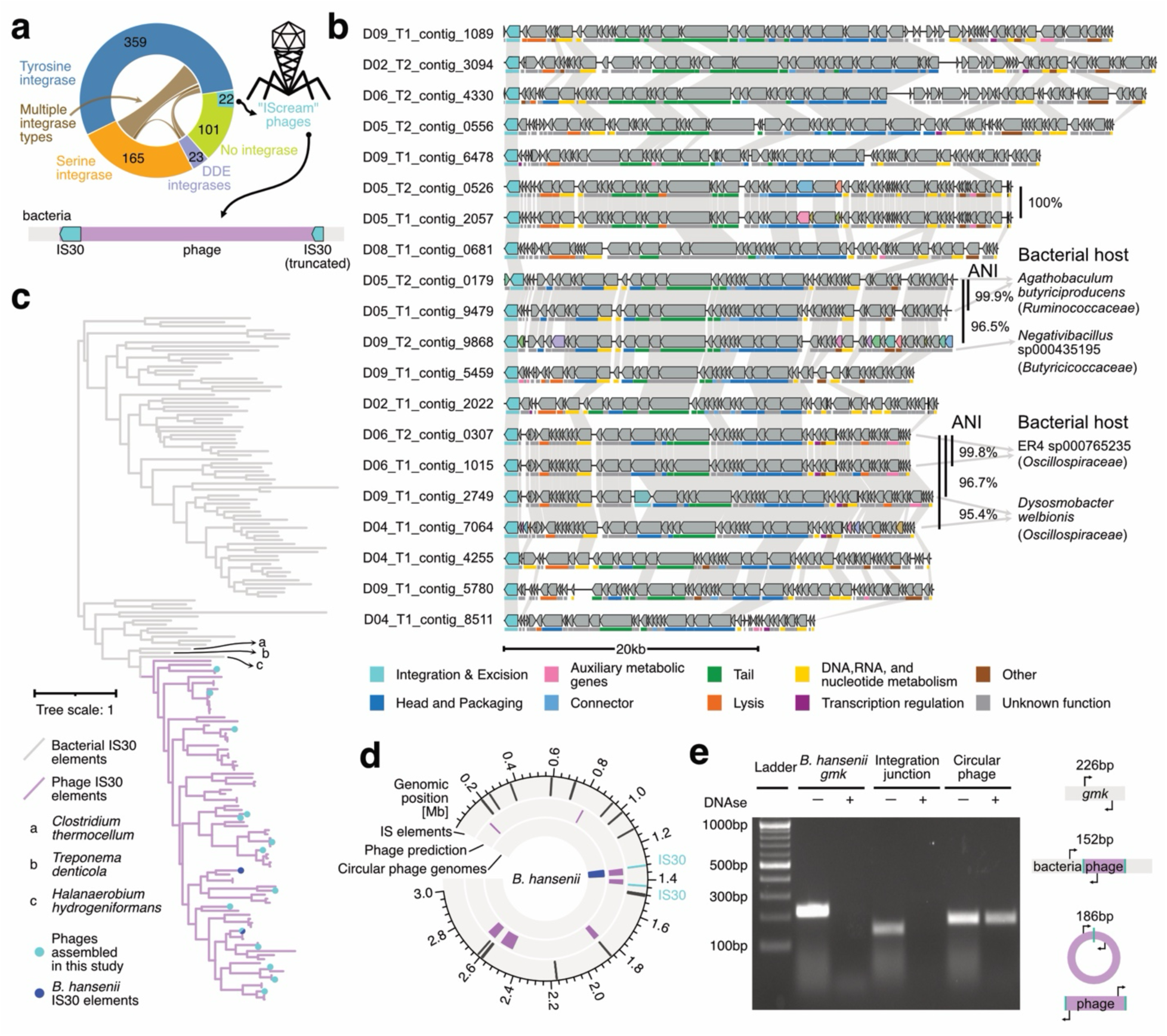
Description of the novel IScream phage group. **A)** Donut plot showing the number of phages with different integrase enzymes. Only phages with SV evidence were included. Of phages without annotated integrase, 22 phages contained genes annotated as IS30 transposases at both ends (schematic below). We named these phages ‘IScream phages’. **B)** Genome organization of full-length IScream phages assembled here, visualized with lovis4u. Gene functionality (annotated as colored bars below genes) was inferred with pharokka. Genes are connected across genomes if the predicted proteins have higher than 25% amino acid identity. Two clusters of closely related IScream phages are annotated with their average nucleotide identity (ANI) and the taxonomic classification of their assembled host context on the right side. **C)** Tree showing the relationship between bona fide bacterial IS30 transposases from the ISfinder database (in grey) and IScream phage outward-directed IS30 (pink) identified in the MGV catalogue. All IS30 transposases were clustered at 70% amino acid similarity before multiple sequence alignment and tree construction. Clusters containing an IS30 from any of the IScream phages assembled here are annotated by cyan dots. **D)** Circos plot for the *Blautia hansenii* ATCC 27552 genome. The outer ring indicates the location of IS elements, the middle ring shows predicted integrated phages, and the innermost ring shows the location of circular phage genomes detected by the presence of structural variation. **E)** Image of a 2% agarose gel showing various PCR products from *B. hansenii* culture, with and without DNAse treatment. The primer location and expected product size are annotated schematically on the right side.

Curiously, among 101 phages lacking an identified integrase we found 22 that contained IS30 family elements on both ends of the prophage genome. This organization is reminiscent of composite transposons: a type of mobile genetic element in which two IS elements flank a gene cassette, encoding e.g. for proteins conferring antibiotic resistance^40^. Since IS30 transposases had been shown in synthetic models to be able to function as phage integrases^41^, we hypothesized that these phages are mobilized through the IS30 within their genomes. Other phages, such as Mu, are known to be mobilized via a transposase^39^; here, we describe a novel group of transposable phages that co-opted bacterial IS30 elements for their mobilization. As this new category of phages likely uses IS30 in a similar manner to a site-specific recombinase, and Cre recombinase (from phage P1) is among the best known phage recombinases in the world, we named these phages “IScream phages” (<u>IS Cre</u>-like phage).

To explore the IScream phages in more detail, we first focused on genome organization and observed high synteny across the full-length IS30-bound phage genomes (see **Fig. 4b**). Genes for core phage lifestyle functions such as structural proteins or host lysis seem to be present in all IScream phages (see **SFig. 9**). Interestingly, two highly similar clusters of IScream phages (ANI > 95%) are present across multiple individuals, integrated into different bacterial host contexts. In fact, one of the order-restricted broad host range phage clusters within the class Clostridia was identified as an IScream phage (see **Fig. 4b**, **STable 2**). Since we observed only a small number of IScream phages in our data, we screened the Metagenome Gut Virus catalogue (MGV) and found 1780 potential IS30-bound phages, revealing that these phages are abundant and prevalent gut residents (see **SFig. 9**).

Focusing more on the IS30 transposase proteins potentially used for phage integration, we clustered all IS30 proteins in the IScream phages assembled here. We observed the IS30 proteins directed outwards of the integrated phage genome to be relatively conserved (mean amino acid identity = 52%), while the IS30 proteins on the other side were highly variable in length (183-4175 nucleotides) and typically truncated or fused to other protein domains, for example domains involved in conjugation (TraX) or defense against restriction (DarA). These deprecated IS30 proteins also typically lack a classical IS30 catalytic domain (see **SFig. 10**), suggesting that the outward-directed IS30 functionally catalyzes phage mobilization. This is again similar to composite transposons, as one of the IS elements in typical composite transposons can lose its catalytic activity and decay over time^42^.

To explore the potential evolutionary history of the IS30 domestication by phages, we clustered the full length, outwardly directed IS30 proteins across the phages identified in our dataset, all IS-bound phages in MGV, and all bacterial IS30 elements annotated in the ISfinder database^43^. We found all phage-derived IS30 proteins to form a sub-clade most closely related to the IS30 elements in *Clostridium thermocellum*, *Treponema denticola*, and *Halanaerobium hydrogenformans* (see **Fig. 4c**). We additionally found IS30 proteins on phages in meso-Amercian paleofeces (see ref ^44^) that clustered together with other phage IS30 proteins (see **SFig. 10**). This suggests that the IScream phages in our data as well as from a paleofeces sample resulted from a single domestication event that potentially occurred in a host related to one of the three species mentioned above.

To gather further evidence that the IS-bound phages represent bona fide phages that can form virions, rather than cryptic prophages or other selfish, non-phage elements, we analyzed publicly available virus-like particle (VLP)-enriched sequencing data from neonatal gut samples^45^ using Phanta, a virus-inclusive read-level profiling method^46^. We found higher abundance of some MGV-derived IScream phages in VLP rather than in metagenomic shotgun (metaG) sequencing (see **SFig. 9**), suggesting viral particle production. We additionally screened Clostridia genomes (see **Methods**) and found two phages flanked by IS30 elements in *Blautia hansenii* ATCC 27552 (see **Fig. 4d**). We obtained and cultured this strain of *B. hansenii* and performed long-read sequencing of the overnight culture. Consistent with our previous results showing a low level of induction for integrated prophages, we observed circular phage genomes in our overnight culture, indicating spontaneous induction of the phage (see **Fig. 4d**, **SFig. 10**). We grew four liters of *B. hansenii* culture and attempted to isolate the phage by PEG-precipitation (see **Methods**). Using PCR, we found the expected circular phage genome in the PEG-precipitate (see **Fig. 4e**). This circular version of the genome, but not the integrated version or the bacterial genome, was protected from DNase treatment, consistent with the presence of phage particles in the culture, where the capsid protects the phage genome from DNase exposure (see **Fig. 4e**).

Together, these findings suggest that IScream phages have co-opted bacterial IS30 transposases for phage mobilization while retaining phage activity, highlighting a previously overlooked group of gut-resident phages.

## Discussion

Decades of detailed research on phages have yielded fundamental insights into molecular biology, genetics, and evolution. Many studies have characterized a select set of culturable phages, which have formed the basis for a collection of ‘central tenets’ of phage biology^2,47^. Though valuable, it is unclear how generalizable these core principles are for such a diverse class of genetic entities. Here, we generated a deep long-read longitudinal metagenomic dataset to derive a more complete understanding of prophage-host interactions within the human gut microbiome. Our findings reveal that a) prophages are usually stable within their hosts over a 2-year timeframe, b) lysogens and naive hosts can co-exist within a population at varying proportions, c) phage induction, when observed, occurs at predominantly at low levels, d) phages can infect bacterial hosts of different families, and e) some prophages might have domesticated insertion sequences as opposed to having integrases for integration and excision.

Previous studies investigating human gut virome dynamics, based on short-read VLP sequencing during a period of up to a year or using bulk metagenomic sequencing over shorter time frames (10 days), have reported generally high stability and individuality of the virome^20,27,48^. Using bulk, long-read metagenomic sequencing and a longer sampling period of two years, we observed most prophages to be stably integrated in our dataset, consistent with stability estimates for the virome. Only in rare cases can we infer possible new phage infection events; for example, we observed an identical phage present in two different strains of *A. putredinis* at T1 and T2 from the same individual (see **Fig. 3a**, **SFig. 5**).

For stably integrated prophages, we commonly observed population heterogeneity in terms of integration, where lysogens and naive hosts co-exist in the same community. Additionally, we were able to quantify heterogeneity when exact boundaries of phage genomes were identified from phage movement over time. This analysis underscores the advantages of longitudinal data and suggests that such heterogeneity may be even more common (see **Fig. 2d**). In a small subset of phages, we similarly detected phage induction through read-level evidence. Within this population, we find that low-level induction is predominant as opposed to large burst events, consistent with short-read-based surveys and ecological models^38,49^. In some cases we find examples where lysogens, naive hosts, and induced circular phage genomes are co-existing in the same population (see **SFig. 6**). This could reflect strain level variation that limits the ability of phages to infect the entire host population. Mechanisms such as phase variation of surface structures within populations have been reported to modulate phage susceptibility^10,50^. Because we do not have information about the spatial distribution of induction in the population, it is difficult to know if induction is a sporadic or somehow coordinated event^51^. Taken together, these findings are consistent with a previously proposed ecological model in which phages spread slowly through a susceptible bacterial population^10,52^, potentially constrained by spatial separation^53^. An added benefit of constant, low-level phage induction might be that it can be a successful strategy for prophages to prevent mutation accumulation and deterioration into cryptic prophages^15^.

The co-evolutionary arms race between phages and bacteria has led to an increasingly narrow host range for many phages^29^. Although some phages can infect hosts within the same genus, some are restricted to the species or strain level^54^. Expansion of the known viral world has led to further investigation of host range among phages, typically using CRISPR spacer analysis or host similarity predictions^37^. Findings from these studies have suggested that broader host ranges may exist up to the class or phylum level^3,4,28,55^. However, CRISPR spacers are typically short (20-50bp) and phage genomes may share large regions due to genome mosaicism^25^, which together can lead to false positive host predictions. In our study, we provide direct evidence supporting the existence of phages with broad lysogenic host range, as we find them integrated into bacterial hosts that differ on the family level. To our knowledge this provides the first direct, molecular evidence of such host ranges present in the human gut. While previous studies have demonstrated a broad host range restricted to the *Bacteroides* genus^56,57^, here we find most phages integrated into distinct taxonomic families have hosts of the *Bacteroidaceae* order. It is important to note, though, that lysogenic host range does not necessarily imply productive host range (the ability to produce infective particles from multiple hosts)^29^. There are many potential determinants for broad host range, such as prophage-encoded diversity generating retroelements^56^, inversions^58^, or polyG tracts^59^, although they have not yet been associated with host range of this breadth. Alternatively, potential order-conserved surface receptor domains or alternative infection routes could explain our findings. The ability of these phages to infect multiple hosts that reside in the gut may have implications for phage therapy that is being explored as a promising alternative to antibiotics^60^. While many of these approaches leverage the narrow host range of cultured phages, evaluating host range via isolation and culturing may limit our ability to detect the breadth of host range^61^. The long-read metagenomic analysis we present here allows us to better capture the landscape of the host range of gut resident phages.

Lastly, we describe the novel group of “IScream” phages, which likely use IS30 transposases of bacterial origin for their mobilization machinery. Previous work had suggested that the distinction between site-specific recombination and transposases might not be well defined^41^. Indeed, Kiss *et al.*^41^ created a synthetic system in which they replaced the integrase of phage lambda with an *E. coli* IS30 and observed that the IS30 transposase was sufficient to create a functional phage. However, the use of IS30 as a recombination machinery for natural phages has not been previously described, to our knowledge. Similar to the transposable phage Mu^39^, IS30 transposases use DDE chemistry; in contrast to Mu, however, we have not yet observed lytic transposition in IScream phages. Using comparative genomic information, we propose a potential model for the emergence of IScream phages: In a bacterial cell of the class *Clostridia*, an IS30 element inserted into an integrated prophage or phage plasmid, that either never had or lost its native integrase. A second IS30 insertion then formed a composite transposon, flanking genes coding for key phage functions such as capsid and tail fiber genes. Over time, the enzymatic activity of the ‘phage-facing’ IS30 open reading frame was lost, while the outward-facing IS30 open reading frame was repurposed for phage mobilization. A successful phage emerged from these events and then diversified into a prevalent and abundant group of phages in the human gut. While it is known that bacteria can domesticate the genes of phages^16^, these findings provide an example for how phages might have domesticated originally selfish genetic elements for their own purpose.

Although we are able to expand our understanding of prophage dynamics in the human gut microbiome through long-read sequencing analysis, this work has several limitations. First, our samples are limited to six healthy individuals from a limited geographic area and therefore our findings may not be generalizable to other populations or physiological states. Our longitudinal samples are also restricted to end-point analyses as we do not have more frequent sampling, which may limit our ability to capture finer dynamics. Second, detection of prophages and their hosts is necessarily linked to sequencing depth. Consequently, despite our deep sequencing in this study, we have not yet exhausted the diversity of the microbiome and are unable to detect organisms at very low abundance. Expanding this analysis to more individuals and timepoints could more broadly reveal the extent of novel phage infection and help elucidate what factors contribute to prophage movement. Third, while *de-novo* assembly allows for reference-free investigation of microbial communities, this approach is not free from errors^62^ and potentially collapses population heterogeneity within a sample. Careful analyses of read-level evidence is therefore needed to support assembly-level claims and quantify the presence of mixed populations. Here, wherever possible, we have used direct read alignment-based approaches to orthogonally validate results derived from assemblies. Fourth, the presence of mixed populations compromises our ability to distinguish between genuine movement of phages and selection on standing variation within a bacterial population. Lastly, phage annotation and host taxonomic classification are subject to error by virtue of their prediction algorithms. For example, the comparison of phage prediction to structural variation evidence revealed that phage annotation tools can misidentify the true boundaries of integrated phages by several kilobases (see **SFig. 6**).

Integrated prophages play a fundamental role in the complex ecosystem of the human gut. They are prevalent and abundant entities that outnumber lytic phages and represent untapped potential for molecular tool development or therapeutic interventions beyond antibiotics. In this article, we reveal key aspects of prophage biology through long-read sequencing, highlighting phage integration dynamics, population heterogeneity, host range, and a novel group of IS-bound phages. We anticipate that future studies will further elucidate the impact of phage-mediated horizontal gene transfer in the gut^17^, characterize the specificity of phage integrases for biotechnological applications, and continue to explore how the evolutionary flux between phages and bacteria may lead to genomic innovation.

## Methods

### Study population

For timepoint 1, stool samples were collected from 6 healthy adult volunteers living in the Bay Area, California, United States. Human subjects research was approved by our institutional internal review board (Stanford IRB 42043; principal investigator: ASB) and informed consent was obtained from all participants. For timepoint 2, the same individuals were re-contacted for a second stool donation, two years after the initial one.

The samples for timepoint 1 were included in the publication of Maghini, Dvorak *et al.*^63^ and short-read metagenomic sequencing reads are available at the National Center for Biotechnology Information’s Sequence Read Archive (SRA) under the identifier PRJNA940499.

### Sample collection and processing

Stool samples were collected without a preservative and stored at -80°C. All DNA extractions were performed using the QIAamp PowerFecal Pro DNA Kit (Qiagen, 51804). DNA extractions were performed according to the manufacturer’s instructions, with the exception of using the EZ-Vac Vacuum Manifold (Zymo Research) instead of centrifugation. DNA concentration was measured using a Qubit 3.0 fluorometer (Thermo Fisher Scientific) with the dsDNA High Sensitivity kit.

### Metagenomic short-read sequencing

Since the samples for T1 had already been sequenced previously, we generated new libraries only for samples from T2. Metagenomic sequencing libraries were pooled and 2x150 base pair reads were generated using the NovaSeq 6000 platform (Illumina; Cat. No. 20012850), to a final depth of 6 Gb per sample.

### Metagenomic long-read sequencing

All samples from both timepoints underwent long-read metagenomic sequencing using the Oxford Nanopore Technology (ONT) platform. DNA fragment distribution was assessed using an Agilent TapeStation (Agilent, Cat. No. G2992AA). Samples with apparent fragmentation were cleaned up using a bead-based protocol before library preparation^64^.

Libraries were prepared with the Native Barcoding Kit V24 (ONT, Cat. No. SQK-NBD114.24), using 1000ng of DNA as input. In the pooling step, four samples were combined, resulting in three total libraries. Libraries were loaded onto PromethION R10.4.1 flowcells (ONT, Cat. No. FLO-PRO114M) and sequenced until exhaustion of the flowcell.

### Short-read data processing

For short-read sequencing, all raw reads were processed with the NextFlow pipeline available under https://github.com/bhattlab/bhattlab_workflows_nf, using NextFlow v22.10.5 30. In short, reads were deduplicated with HTStream SuperDeduper v1.3.3 and low-quality bases were trimmed with TrimGalore v0.6.7. Then, reads were mapped against the human genome (hg38) using bwa v0.7.17 31 and all matching reads were discarded. Since the samples from T1 had been sequenced very deeply and with highly variable library sizes (ranging from 8Gbases to 15Gbases), we downsampled preprocessed reads of T1 samples to a final library size randomly sampled from the distribution of final library sizes of T2 samples after preprocessing.

For each sample, metagenomic assembly was performed with metahit v.1.2.9 and genes were predicted with bakta v1.8.2. Assemblies were binned into draft genomes using MetaBAT v.2.5, CONCOCT v.1.1.0, and MaxBin v.2.2.7, followed by bin consolidation using DAStool v1.1.6. Bin quality was assessed using CheckM v1.2.2 and taxonomic classification was performed using GTDB-tk v2.3.0 using the GTDB r214 database.

### Long-read data processing

For long-read sequencing, pod5 files were basecalled and de-multiplexed with dorado v0.5.3,, using the ‘super-high accuracy’ model v3.4.0, to create the final set of fastq files. Read quality and length distribution was assessed using NanoPlot v1.41.6 before and after removal of human reads (reads mapping against the human genome (v38) using minimap2 v2.26-r1175). Metagenomic assembly was performed with meta-flye v2.9.2-b1786, using the nano-hq flag to use only reads of quality Q20 or higher for initial assembly. Binning and taxonomic classification of bins were performed as described for short reads. The workflows for long-read data processing are also available under https://github.com/bhattlab/bhattlab_workflows_nf.

In order to better compare short and long reads, long reads were subsampled to the same mean depth as the short reads using custom scripts, randomly selecting read ids from the processed reads up to a specified amount of total sequencing. Seqtk subseq v1.4-r130 was used to subsample the original fastq files for the selected read ids. Read quality and length distribution was assessed using NanoPlot v1.41.6. Reads were assembled and binned using the same workflow as above.

### Phage prediction

To predict phages, we applied geNomad v1.7.6^24^, VIBRANT v1.2.1^22^, and virsorter2 v2.2.4^23^ to the short and long-read assemblies for each metagenomic sample. For each predicted phage, we used CheckV v1.0.1 ^36^ to assess their quality. VIBRANT, geNomad, and CheckV are included in the NextFlow project available under https://github.com/bhattlab/bhattlab_workflows_nf. Virsorter2 was run separately since it relies on snakemake (v5.26.0) for execution. All predicted phages (both full phage contigs and predicted prophages) were collated from all three tools.

### Comparison across short and long-read sequencing

To compare the phage predictions across short and long-read sequencing, we mapped short-read contigs against the (subsampled) long-read assembly using blast v2.2.31+, filtering alignments for identity (99%) and query coverage (90%). To determine if an integrated long-read phage was fully covered by short-read contigs, we required a single short-read contig to align to at least 95% of the phage region. To determine the short-read coverage of integrated phages, we mapped the short reads to the long-read assembly with bowtie2 v2.5.4 and calculated the per-base coverage with samtools v1.21.

To compare short and long-read phage predictions in more detail, we adapted the segment overlap metric (originally developed to measure overlaps between predicted biosynthetic gene clusters^65^) to measure which fraction of long-read phages is covered by predicted short-read phages and vice versa (see **SFig. 4**). This metric calculates recall by considering each long-read phage prediction as a positive instance. If a given long-read phage is covered (up to a variable cutoff of x%) by an alignment of a or more short-read contigs predicted to be phage, we annotate it as true positive, otherwise as false negative. Recall is then defined as true positives over the sum of all positives, in effect quantifying the fraction of long-read phages that are found by short-read sequencing as well. Similarly, precision is calculated based on the short-read phages, with short read phages overlapping (to at least x%) a long-read phage to be considered true positives, whereas those not overlapping a long-read phage to be considered false positives. Precision is then calculated as true positives over the sum of true and false positives, quantifying the fraction of short-read phages found by long-read sequencing as well.

### Clustering of phages across timepoints

To cluster prophages across timepoints within each individual, we used blast v2.2.31+ to map all phages of T1 and T2 against each other and then computed genome identity and coverage with the CheckV companion scripts ^36^. Additionally, we extracted 10kb regions on either side of integrated prophages and performed the same analyses. We then consolidated the clustering by comparing the identity and coverage across clusters created from the phage identity and coverage file only (see **SFig. 5**), using high identity (99%) and genome coverage (90%) cutoffs to identify the same phage across timepoints. Since we observed large structural variations between phages integrated into the same host context, we relaxed the genome coverage cutoff for clustering when the host context was identical. Similarly, as phages were sometimes identified at the beginning or end of contigs, we reduced the host-region coverage cutoff to be 40% to include cases where only one side of the host region was found because of fragmented assemblies.

### Identification of structural variation that are overlapping predicted phage regions

To find structural variations in our data, we mapped the reads of each timepoint against the assembly of the other timepoint within an individual, using NGMLR v0.2.7^66^. Then, bam files were sorted with samtools v1.9 and structural variants were called using Sniffles2 v2.2^67^.

### Assignment of host taxonomy for integrated prophages

We used iPHoP v1.3.3^37^ to predict hosts for all phages, using default parameters and database version iPHoP_db_Aug23_rw. For integrated long-read phages, we annotated their hosts by classifying each annotated gene on a given contig using the mmseqs-taxonomy module from mmseqs2 v14.7e284^68^. This module provides a taxonomic annotation based on the GTDB v214.1^69^ database for each gene. For each contig, we then combined the annotations for all genes not located in predicted phage regions at each taxonomic level. Annotations were accepted if more than 50% of genes agreed, disregarding genes without taxonomic annotation.

iPHoP, mmseqș2-based, and binning taxonomic assignments were evaluated for consensus at each taxonomic level (see **SFig 4**). To determine a subset of integrated phages for which we had high confidence host assignment, we filtered our results for agreement between high-quality bins (>90% completeness and <5% contamination, see ref^70^) and mmseqs2-based assignment down to the family level.

### Clustering of phages on species-level for host-range analysis

To evaluate host-range within phage species, we clustered all geNomad annotated phages in all samples at a minimum of 95% ANI and 80% alignment fraction using CheckV v1.0.1 ^36^ supporting scripts. We specified a minimum 80% query and target coverage (--min_tcov 80 --min_qcov 80), as we expect phages to be assembled more contiguously by long reads. The resulting cluster membership information was merged with the host annotations from our mmseqs2-taxonomy approach and bin taxonomy, described above.

### Gene content for integrated prophages

To assess gene content across all integrated prophages with structural variant evidence, we used Pharokka v1.7.3 against the Pharokka v1.4.0 database ^71^ with the --meta, --skip-mash, and --split flags.

### Synteny and gene content visualization

Visualization of all gene content and synteny was done using LoVis4u v0.1.4.1 ^72^. Two of the 22 IScream phages were smaller than 10kb and therefore were removed from this analysis. For IScream phage synteny visualization, gff files of the identified full length IScream phages, generated by Pharokka annotation (described above), were used as input for LoVis4u visualization with default configuration. For host context visualization, a custom script was used to parse through gff files from bakta to visualize specified windows and to convert them to a format compatible with LoVis4u. The reformatted gff3 files were used as input for LoVis4u visualization using an updated configuration file specifying mmseqs_min_seq_id = 0.8 from the 0.35 default value.

### Identification of potential integrase enzymes

To identify potential integrase enzymes in our assemblies, we used a set of Pfam Hidden-Markov-Models described in an earlier exploration of mobile genetic elements^73^: PFPF07508 for large serine recombinases, PF00239 for small serine recombinases, PF00589 for tyrosine recombinases, and PF00665, PF13333, and PF13683 for DDE recombinases. We used the hmmsearch command (with the `—cut_ga` flag) from HMMER v3.4 ^74^ against all predicted proteins. We then annotated each phage by counting which type of potential integrase was present within the phage boundaries, using only phages with evidence from structural variations.

### Identification of IScream phages in MGV and Clostridia genomes

To explore the prevalence and abundance of IScream phages, we searched for potential IScream phages in public datasets. We ran ISEscan on all viral genome assemblies from MGV, and identified genomes that contained one IS30 element starting less than 1kb from the beginning of the assembly and one IS30 element ending less than 1kb from the end of the assembly.

We next searched for existing bacterial isolates containing integrated IScream phages. We downloaded all 2160 bacterial genome assemblies annotated to have assemblies at complete or chromosomal level, and annotated as belonging to the Clostridia class, using the NCBI Datasets tool. We ran ISEscan and geNomad on all assemblies, and identified genomes that contained a geNomad annotated prophage region with one IS30 element starting less than 1kb from the beginning of the prophage region and one IS30 element ending less than 1kb from the end of the prophage region, which included Blautia hansenii ATCC 27552 (see **STable 3** for a complete list of potential IScream phages).

Additionally, we analyzed the species-level taxonomic profiles for two public datasets, generated with phanta v1.0 ^46^, which included a viral database built on representative genomes from MGV (see the original phanta publication for details about data processing). Each phage species from this database was classified as potential IScream phage, if a genome contained in this species bin was identified to be an IScream phage.

The dataset from Liang *et al.* ^45^ included bulk metagenomic sequencing and virus-like particle (VLP)-enriched sequencing of infants. In this dataset, we quantified the relative taxonomic abundance of IScream-containing phage species in paired bulk and VLP-sequencing in order to identify potential particle formation of IScream phages. Lastly, the non-cancer controls from the dataset from Yachida *et al.* ^75^ was used to calculate prevalence and mean abundance of phage species in healthy individuals.

### Clustering of IS30 elements

To cluster IS30 elements across IScream phages, we used the ete3 toolkit v3.1.3 ^76^. First, we build a tree for all IS30 proteins on all IScream phages using the ‘standard_fasttre’ workflow from ete3, consisting of a multiple sequence alignment with Clustal Omega v1.2.4 and tree construction using FastTree v2.1.8 (see **SFig. 10**). Since this analysis showed a clear separation between outward and inward-directed IS30 proteins, we afterwards focused only on the outward-directed IS30. To gain insights into the evolutionary history of IScream phages, we extracted all outward-directed IS30 proteins from our IScream phages, all potential MGV IScream phages, and all bona-fide bacterial IS30 elements from the ISfinder database. For the MGV phages, we filtered the outward directed IS30 proteins to be longer than 600 and shorter than 2200 nucleotides. All proteins were clustered at 70% amino acid similarity over 80% of the alignment, using mmseqs v14.7e284^77^. Then, a tree was constructed on the cluster representatives using the ‘standard_fasttre’ workflow in ete3, modified to trim positions in the multiple sequence alignment with more than 90% gaps using trimAl v1.4.rev6.

### Identification of potential IScream IS30 elements in ancient stool metagenomes

To identify potential IScream phages in ancient stool samples, we downloaded the raw data from Wibowo *et al.*^44^, containing sequencing of 1000-2000 year old desiccated paleofecal samples from the southwestern USA and Mexico. Raw reads were processed and assembled as described above and phages were identified with geNomad and IS elements with ISEscan. For all contigs that contained IS30 elements within predicted phages, we assessed their DNA damage level with DamageProfiler v1.1 ^78^. To prevent inclusion of modern IScream phage IS30 proteins, we filtered the phage-overlapping IS30 proteins for being on contigs recruiting more than 1000 reads and showing an estimated 5‘C’T damage level or an estimated 3‘G’A damage level over 1%, resulting in six potential IScream IS30 open reading frames. Note that these filtering steps are rather specific, since many more IS30 proteins are identified on shorter contigs which are not predicted to be of phage origin. The six ancient IS30 proteins were added to the clustered IS30 proteins from MGV, IScream phages, and ISfinder and the tree was recomputed as described above.

### Culturing of the B. hansenii IScream phage

*Blautia Hansenii (*ATCC 27552) was grown anaerobically (90% nitrogen, 5% carbon dioxide, 5% hydrogen) in an anaerobic chamber (Sheldon Manufacturing) in hemin (5 μg ml^−1^), l-cysteine (1 mg ml^−1^), and sodium bicarbonate (0.2%) supplemented Brain Heart Infusion (Sigma) media (BHIS).

*B. hansenii* was grown overnight in pre-reduced BHIS. The culture was pelleted by centrifugation (Eppendorf, 5920R) at 4000 g for 15 minutes and DNA was extracted using the Qiagen DNeasy Blood and Tissue kit (Qiagen, Cat. No. 69504) following manufacturers instructions for Gram positive bacteria. Bacterial genome sequencing was performed by Plasmidsaurus using Oxford Nanopore Technology with custom analysis and annotation.

### PEG precipitation of B. hansenii phage particles

*B. hansenii* was grown in 4 liters of BHIS overnight. Supernatant from the culture was harvested by centrifugation (Eppendorf, 5920R) at 4198 g for 15 minutes at 4C. Sodium chloride was added to the supernatant to a final concentration of 5M and PEG-8000 was added to a final concentration of 10%. This solution was stirred for 30 minutes at 4C to dissolve and then left at 4C with no agitation overnight. PEG-precipitated phage particles were harvested by centrifugation at 10,000 x g for 10 minutes at 4C. Supernatant was removed and the resulting pellets were allowed to air dry for 3-5 minutes inverted. The pellets were combined and resuspended in a total of 10mL of SM buffer (100 mM NaCl, 8mM MgSO4*7H2O, 50mM Tris HCl pH 7.5). An equal volume of chloroform was added, mixing by inverting, and centrifuged at 12,000 x g for 10 minutes at 4C. The aqueous phase was collected and chloroform treatment was repeated.

### PCR validation of phage particles from B. hansenii IScream phage

PCR primers were designed using NCBI Primer Blast under default settings. PCR product size was targeted to be between 150 and 250 bp. PCR primers were designed to target 1) *B. hansenii* gmk gene, 2) an internal region of the IScream phage, 3) the upstream junction site of the integrated phage and bacterial chromosome, and 4) to span the junction of circularization of the IScream phage region (see **Fig 4e**; primers listed in **STable 4**). PCR reactions were performed on PEG precipitated samples directly with and without DNase treatment. For the DNase treated samples, 50uL of PEG Prep was treated with 5ul 10X TurboDNase Buffer, 1.13uL TurboDNase (Invitrogen™ AM2239), and 1uL RNase (1mg/mL Invitrogen™ AM2270), incubated at 37C for 1 hour and then heat inactivated at 70C for 10 minutes.

PCR reactions were performed using Q5 high fidelity polymerase (67 °C annealing temp, 30 s annealing time, and 5 s extension time at 72°C). PCR reactions were run on a 2% agarose gel.

### Statistical analysis and visualization

All statistical tests were performed in R v4.2.2. Data visualization was performed using ggplot2 v3.5.1, which is part of the tidyverse v2.0.0 suite of tools ^79^.

## Data availability

The raw sequencing data for all samples sequenced in this study are available from the European Nucleotide Archive under the study identifier <u>PRJEB88320</u>. The short read sequencing data from T1 had been included in the earlier publication by Maghini, Dvorak *et al.* 2024 and are available under the identifier <u>PRJNA940499</u>. Data for the Metagenome Gut Virus catalogue (MGV) from the publication by Almeida *et al.* 2020 is available under https://portal.nersc.gov/MGV/. The raw sequencing data of ancient metagenomic samples from the publication by Wibowo *et al.* 2021 are available under <u>PRJNA561510</u>. The raw data for the phanta-profiled datasets from Liang *et al.* 2020 is available <u>PRJNA524703</u> and under <u>PRJDB4176</u> for the study from Yachida *et al.* 2019.

## Code availability

Source code for analysis and figure generation is publicly available at Zenodo under https://doi.org/10.5281/zenodo.15192469 and on GitHub under https://github.com/bhattlab/long_read_benchmark.

## Acknowledgement

We would like to thank all the members of the Bhatt lab for enriching discussions and invaluable feedback, especially Dylan Maghini and Yishay Pinto. We would also like to thank members of the Ashley, Altemose and Good labs at Stanford university for help with Nanopore sequencing and inspiring discussions. We would like to thank Xianfeng Zeng from the Fischbach lab for providing the *Blautia hansenii* ATCC 27552 sample.

ASB is supported by the Paul Allen Distinguished Investigator Award and the Bhatt lab is supported by a Stand Up 2 Cancer Grant and NIH R01AI148623 and R01AI143757. JW is a Damon Runyon Quantitative Biology Fellow supported by the Damon Runyon Cancer Research Foundation (DRQ-22-24). ASH and DTS acknowledge support from the National Science Foundation Graduate Research Fellowship Program (DGE-1656518). DTS and MD are supported by the Cell and Molecular Biology Training Grant (T32 GM007276). RBC is supported by the A.P. Giannini Foundation.

## Author contributions

Sample collection and processing: MD

Long-read sequencing: MD, ASH, RBC, JW

Blautia experiments: ASH, DC, RBC, DTS

Data analysis: ASH, JW, DC, NE

Conceptualization: JW, ASH, ASB

Supervision: JW, ASB

JW, ASH, and ASB wrote the manuscript with input from all authors. All authors read and approved the final manuscript.

## Supplemental information

## Supplementary Tables

STable 1: Sequencing stats

STable 2: Broad host range examples

STable 3: Other IScream phages in Clostridial genomes

STable 4: Blautia primer sequences

## Supplementary Figures

SFig. 1: Sequencing stats and quality

SFig. 2: Comparison to other phage prediction tools

SFig. 3: overlap metrics for short and long-read sequencing

SFig. 4: Host identification comparisons

SFig. 5: Methods for clustering phages

SFig. 6: Detection of structural variations with sniffles

SFig. 7: Alistpes bin comparison

SFig. 8: Synteny and coverage plot for examples with broad host range

SFig. 9: IScream phage gene content and MGV evidence

SFig. 10: IScream phage IS30 tree & similarity, ancient genome IS30 elements

