## Supplementary Figures for "Discovering Broader Host Ranges and an IS-bound Prophage Class Through Long-Read Metagenomics"

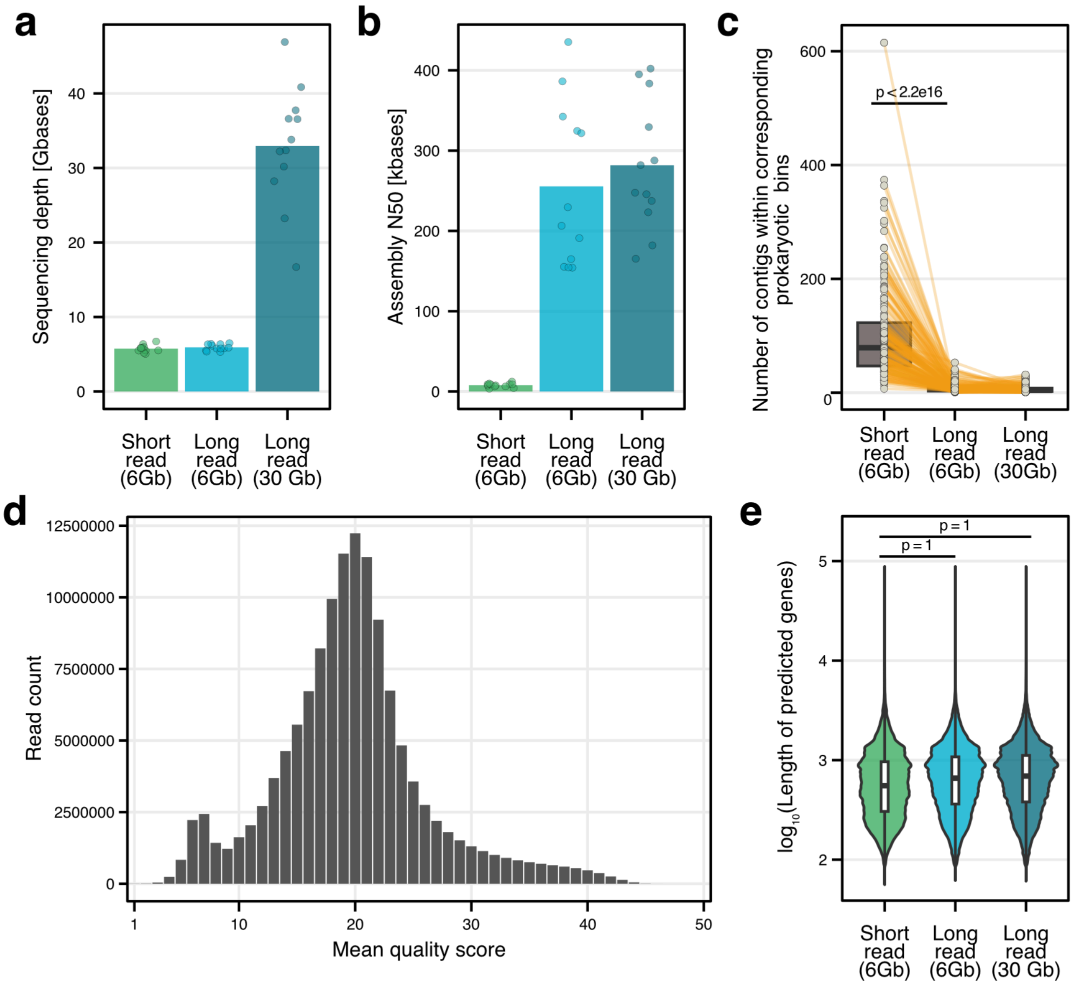


**Supplementary Figure 1: Comparisons between short-read and long-read sequencing. a)** Mean sequencing depth of Illumina short read sequencing, subsampled ONT long-read sequencing, and deep ONT long-read sequencing across all samples are shown as bars. Points indicate individual samples. **b)** Mean assembly n50 across different sequencing types and depth as bars while points indicate individual samples. **c)** Comparison of the number of contigs present in assembled bins that were shared between the three sequencing types: Illumina short reads (SR) at a depth ~6Gb, subsampled ONT reads (LR) to ~6Gb, and deep ONT reads (DLR) at ~30Gb, represented as a box plot. Boxplots show the interquartile ranges (IQRs) as boxes, with the median as a black horizontal line, whiskers extending up to the most extreme points within 1.5-fold IQR, and all data points are indicated as dots. Corresponding bins were determined by matching bin taxonomic assignment across sequencing approaches. Orange lines track corresponding bins across the sequencing types. A paired Wilcoxon signed rank test was used to compare the number of contigs per bin from short reads at ~6Gb depth and long reads at 6Gb depth, finding that there is a significant difference (n = 183, p-value < 2.2e-16). **d)** Histogram of mean read quality score from deep long read sequencing samples. **e)** Comparison of the length of predicted genes from assemblies of each sequencing type. A one-sided Wilcoxon rank sum test was used to compare the median length of predicted genes between Illumina short reads and ONT long reads subsampled to the same depth. Results showed no significant difference (SR n = 3457575, LR n = 3408004, p = 1). Similarly, one-sided Wilcoxon rank sum test was used to compare the short read assembly with the deep ONT assembly, similarly showing no significant difference (SR n = 3457575, DLR n = 6671897, p = 1) The Wilcoxon test indicated that the median gene lengths from short read assembly were not significantly greater than median gene lengths from ONT assemblies.  Boxplots show the interquartile ranges (IQRs)

as boxes, with the median as a black horizontal line and the whiskers extending up to the most extreme points within 1.5-fold IQR. Violin plots show overall sample distribution.

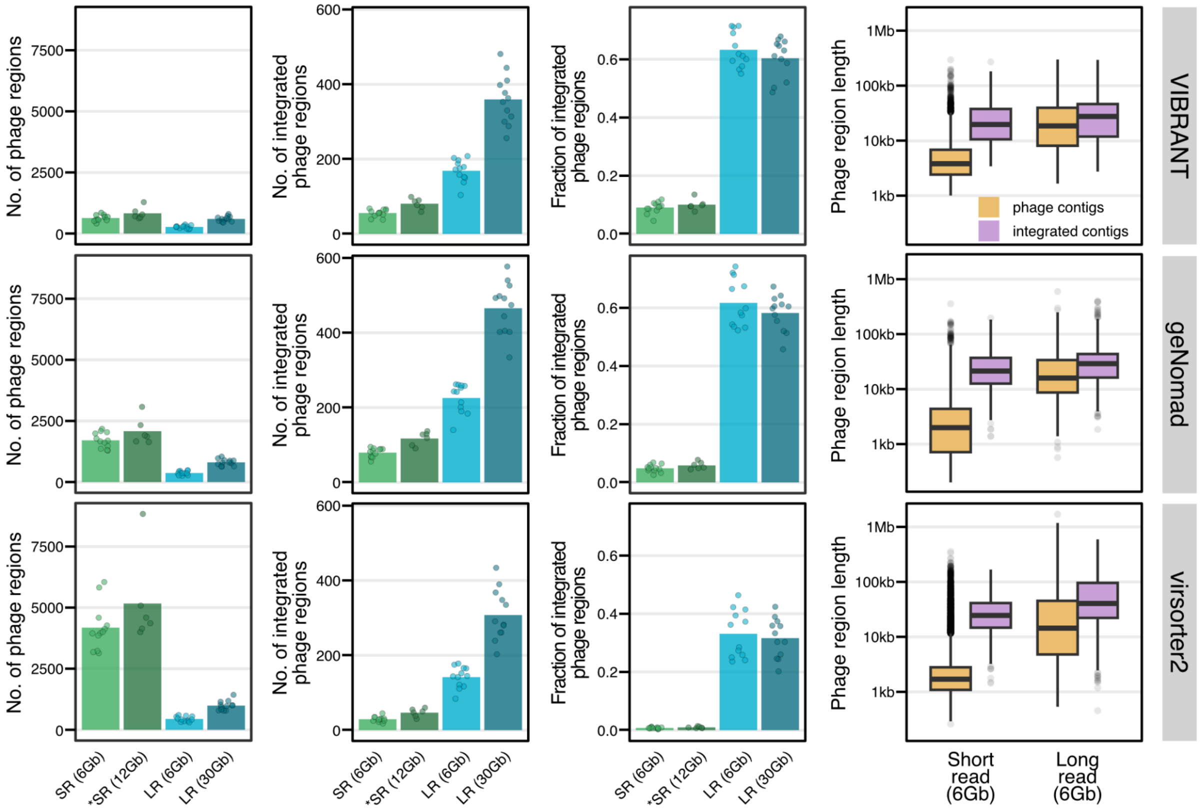


**Supplementary Figure 2: Comparisons between different phage annotation tools** For each phage annotation tool (VIBRANT, geNomad, and virsorter2), the average total number of phages annotated, average number of integrated phages annotated, and the average fraction of identified phages that were integrated are presented for each sequencing type and depth (Illumina short reads ~6Gb, Illumina short reads ~12Gb (T1 only, n = 6)*, ONT long reads ~6Gb, and ONT long reads ~30Gb). Phage length was compared between entire phage contigs (orange) and integrated phages (purple) for Illumina short reads at ~6Gb and subsampled ONT long reads at ~6Gb, indicating that short-read sequencing resulted in more fragmented (and therefore shorter) phage genomes. Boxplots show the interquartile ranges (IQRs) as boxes, with the median as a black horizontal line, whiskers extending up to the most extreme points within 1.5-fold IQR, and outliers indicated as dots.

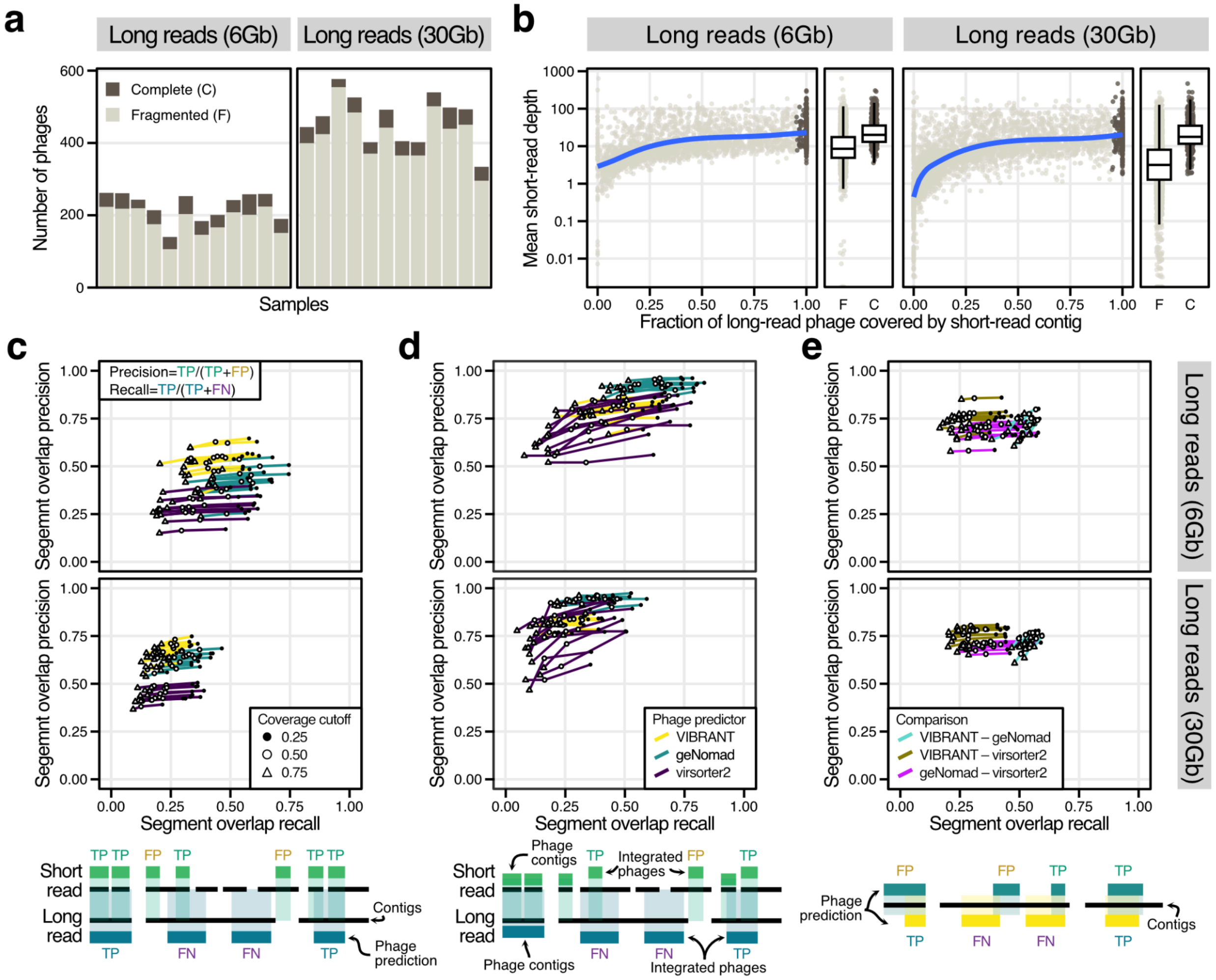

**Supplementary Figure 3: Short-read phage fragmentation and segment overlap analysis**

**a)** Barplot showing the number of integrated phages per sample for downsampled (~6Gb) and deep (~30Gb) long-read sequencing. The fill color indicates if the same phage was found to be fragmented in the short-read assembly or found to be completely covered by a single short-read contig. **b)** Short-read depth of all integrated prophages identified with geNomad in the long-read assemblies. Short reads were mapped to the long-read assemblies and the average depth for all regions identified to be phages was calculated. Dots indicate if the phage was found to be fully covered or fragmented when mapping short-read contigs to the long-read assemblies. The right-side panel for each plot shows the average short-read depth as boxplots. Boxplots show the interquartile ranges (IQRs) as boxes, with the median as a black horizontal line and the whiskers extending up to the most extreme points within 1.5-fold IQR. **c)** Segment overlap metric (see **Methods** and ref Carroll et al.) calculated between short-read phages and long-read phages. Across all predictors, only about 20% of long-read phages were found to be sufficiently covered by short-read phages (segment overlap recall). This fraction increased with a more lenient cutoff for how much of the long-read phage had to be covered to be considered a true positive, indicating that only a small part of the long-read phage is covered by short-read phage contigs. Similarly, the segment overlap precision (how many short-read phages overlapped long-read phages) was higher for all phage predictors, but decreases with the overall number of phages predicted to be present. Especially for virsorter2, only about 35% of the circa 4000 predicted phages per sample overlapped corresponding long-read phage predictions, indicating that virsorter2 predicts many contigs to be phage which it does not predict as phage when assembled in context. See schematic at the bottom of the figure panel for a visual representation of mapping of short-read phages to long-read phages. **d)** Same as panel **c**, but only for integrated phages (disregarding phage contigs). In this analysis, segment overlap precision is ~80%, indicating that most of phages identified to be integrated prophages in short-read assemblies are similarly identified in the long-read assemblies, whereas the coverage of integrated phages in the long-read assemblies remains relatively low (~20%). See schematic at the bottom of the figure panel for a visual representation of mapping of integrated short-read phages to long-read phages. **e)** Segment overlap metric when comparing across phage predictors, in the long-read assemblies only. See schematic at the bottom of the figure panel for a visual representation for how segment overlap and recall were calculated when comparing predictors.

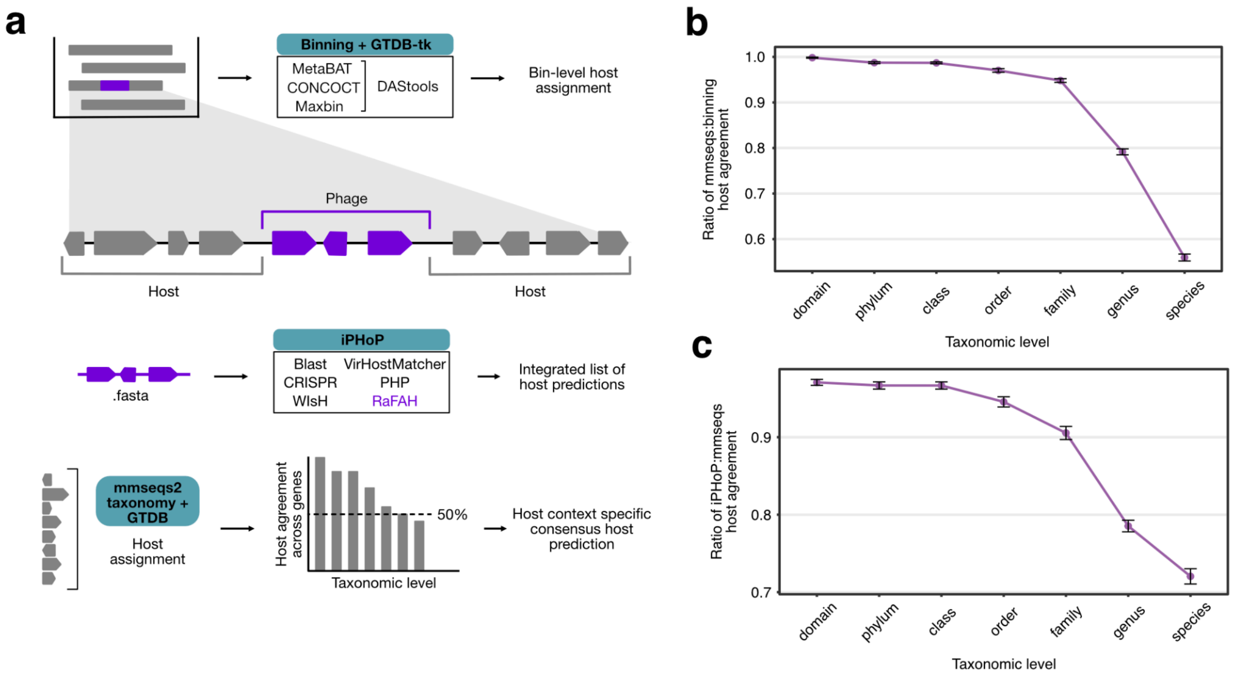


**Supplementary Figure 4: Host taxonomy assignment schematic and benchmarking**
**a)** Schematic representation of the different approaches to assigning hosts to integrated phages. From the long-read assemblies, we assign bacterial hosts for a given prophages by comparing the bin taxonomic assignment from GTDB-tk (bin membership of the contig containing the integrated prophage, see **Methods**) and gene-level annotations from mmseqs-taxonomy (see **Methods**). For this approach, all genes in host regions of the same contig as the integrated prophage are annotated against the GTDB database and a consensus taxonomic assignment is generated by majority rule. The current gold standard for taxonomic prediction of phage hosts is the prediction tool iPHoP, which integrates the predictions from several different approaches. Host prediction via iPHoP has traditionally been necessary, since phages are often assembled as fragments or without host context with short-read sequencing. For each phage genome, iPHoP generates an integrated list of host predictions. **b)** Mean ratio of the host agreement between the mmseq2-taxonomy based approach and binning taxonomic assignment for integrated prophages across all samples at each taxonomic level. Error bars indicate the standard error across all samples. **c)** Mean ratio of the host agreement between the mmseq2-taxonomy based approach and iPHoP taxonomic assignment for integrated prophages across all samples at each taxonomic level. Since iPHoP outputs a list of possible hosts, here we looked for agreement at each taxonomic level from any potential host. Error bars indicate the standard error across all samples.

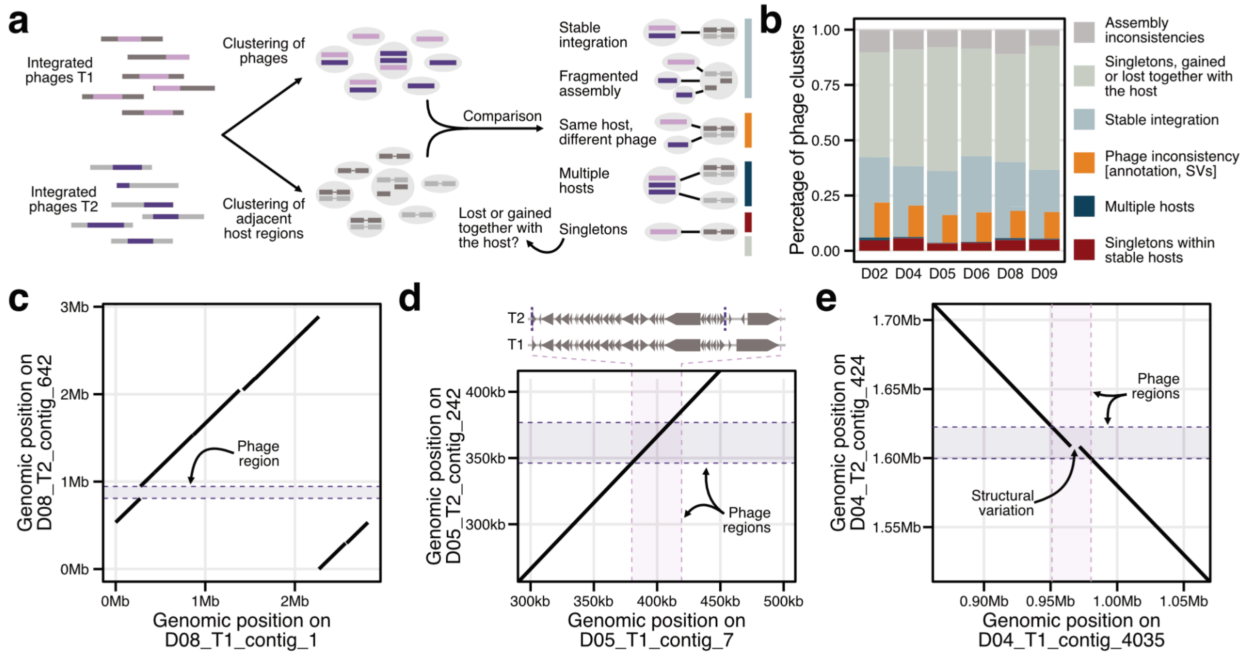


**Supplementary Figure 5: Clustering of prophages within individuals.**
**a)** Schematic showing the clustering and classification of integrated phages within individuals. In short, all integrated phages and their adjacent host regions were clustered separately using the CheckV companion scripts for genome identity and coverage calculation based on blast mappings. Clustering was done with high identity (99%) and genome coverage (90%) cutoffs. Then, clusters were refined by comparing between phage and host clusters. **b)** Detailed classification of phage clusters per individual (see **Fig. 2** in the main text). Clusters with phage inconsistencies were classified as stable integration, since the inconsistencies were due to differences in annotation or large structural variation within phage genomes, yielding genome coverage values below our cutoff for clustering. **c)** Dotplot comparing two contigs within individual D08, showing the gain of a 137kb integrated phage into an otherwise stable host. The phage region (representing a dynamic phage) is indicated by a shaded pink area. **d)** Dotplot comparing two contigs within individual D05, showing a cluster classified as ‘phage inconsistency’ due to variations in phage annotation. The phage regions are indicated by shaded pink areas. While both contigs align perfectly, the region annotated as phage in both timepoints do not overlap over more than 90%, resulting in disparate clustering outcomes. The reason for this inconsistency is a single nucleotide polymorphism changing the predicted genes in T2 (annotated above). **e)** Dotplot comparing two contigs within individual D04, showing a cluster classified as ‘phage inconsistency’ due to a large structural variation (deletion in T2). The phage regions are indicated by shaded pink areas. The gap in the alignment represents a 6.5kb deletion in T2 compared to T1, resulting in disparate clustering outcomes because of respective genome coverage below 90%.

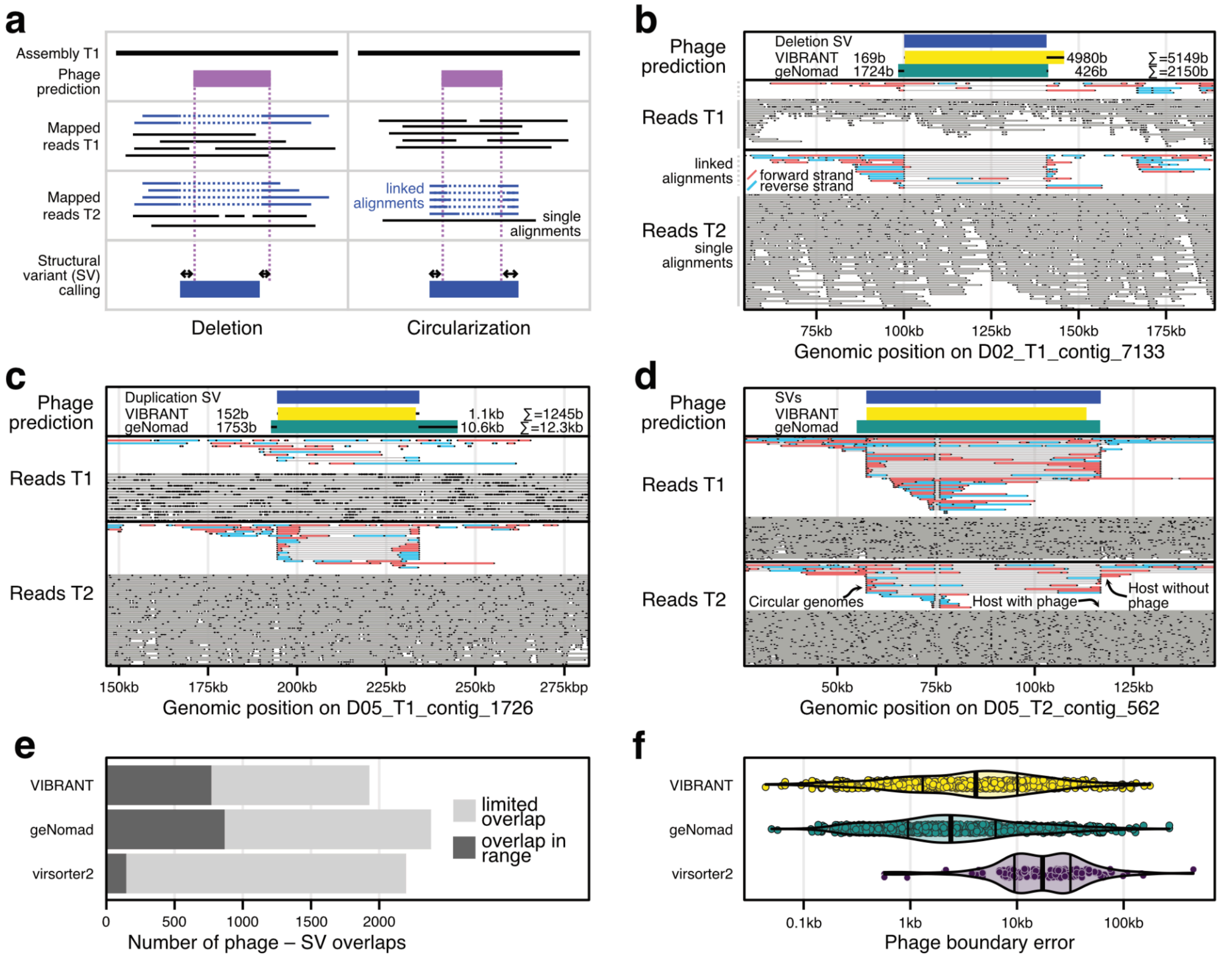


**Supplementary Figure 6: Detection of exact phage boundaries through long-read mapping and structural variation calling.**

**a)** Schematic representation of the identification of exact phage boundaries by structural variant (SV) calling. In short, SVs were identified with Sniffles2 on the basis of read mapping within and across timepoints for the same individual. In these cases, linked read alignments (either supporting a deletion or a duplication, represented by blue lines in the schematic) can be used to identify SVs. Duplication SVs can be interpreted as the presence of circular phage genomes. We considered all phage-SV overlaps that covered at least 50% of both phage and SV to be high-quality phages identified with base-pair accuracy. **b)** Read alignment plot illustrating a phage identified by a deletion SV. Read alignments are separated into linked alignments and single alignments. Each line represents the alignment of a single read with dots showing the start and end of each alignment. Linked alignments are colored by strand and connected by a thin grey line to indicate that they originate from the same read. On the top, the coordinates from phage predictions and the process of boundary evaluation is shown: the absolute difference on either side of the prediction is summed to create the total phage boundary error. **c)** Same plot as **b**, but for a duplication SV (circular phage genome), identified in individual D05. **d)** Same plot as **b**, showing for a phage region present in individual D05 that there is read evidence for the simultaneous presence of circular phage genomes (linked reads supporting a duplication SV), hosts without the integrated phage (linked reads supporting a deletion SV), and hosts with the integrated phage (single alignments) in T2. **e)** Number of phages predicted by different tools that overlapped SVs called by Sniffles2, colored by their overlap being in range (more than 50% of the phage and the SV region) or not within range. Virsorter2 tends to predict very long phages, resulting in many comparisons where less than 50% of the phage was covered by the SV region. **f)** Evaluation of the boundary error (see **bc**) for all phage-SV overlaps within range (see **e**). Each dot represents a phage-SV overlap, the overall distribution is shown as violin, and vertical bars indicate the 25%, 50% (bold) and 75% percentile.

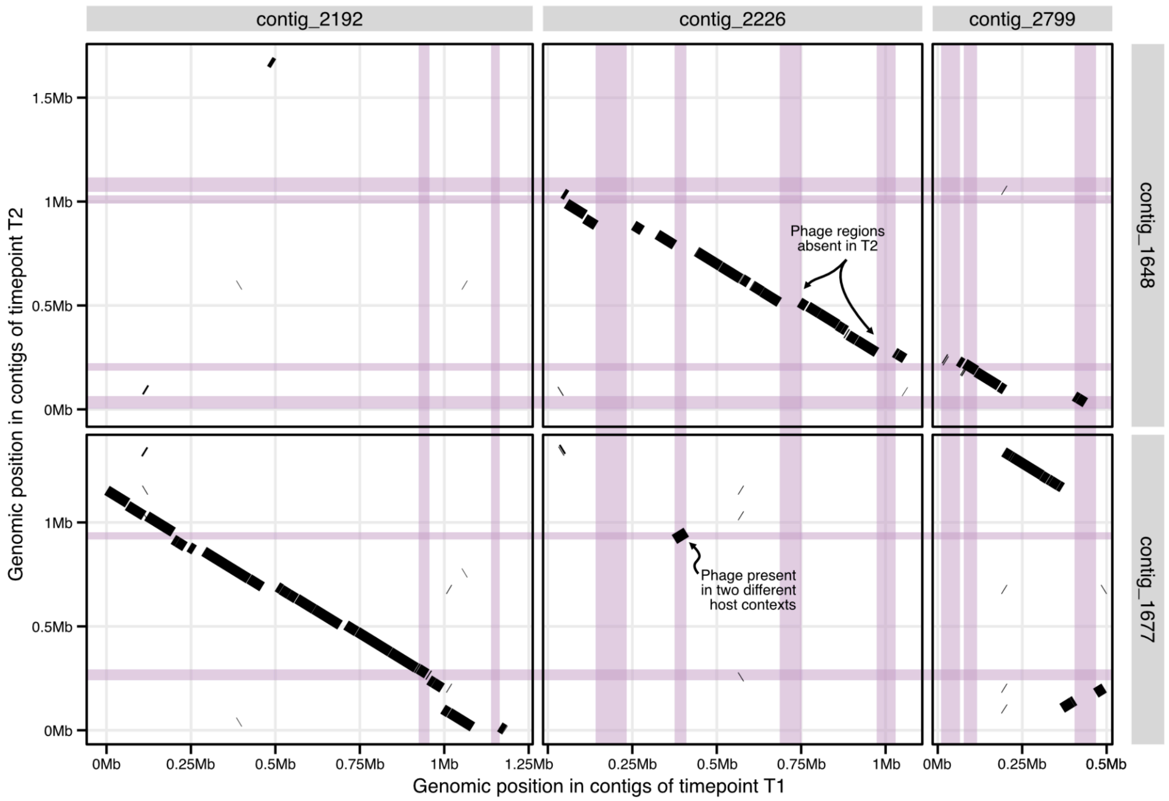


**Supplementary Figure 7: Genome comparison between the two *Alistipes putredinis* metagenome-assembled genomes.**

Dotplot comparing two *Alistipes putredinis* metagenome-assembled genomes in individual D09, generated by nucmer (see **Methods**). Only contigs longer than 10kb were included in this analysis. Annotated phages are shown by shaded pink areas, revealing multiple regions of structural differences between both assemblies due to variations in phage content. The phage region that is present in T1_contig_2226 and T2_contig_1677 is classified as a dynamic phage, since the phage genome between both time points is identical, but the surrounding host regions are not.

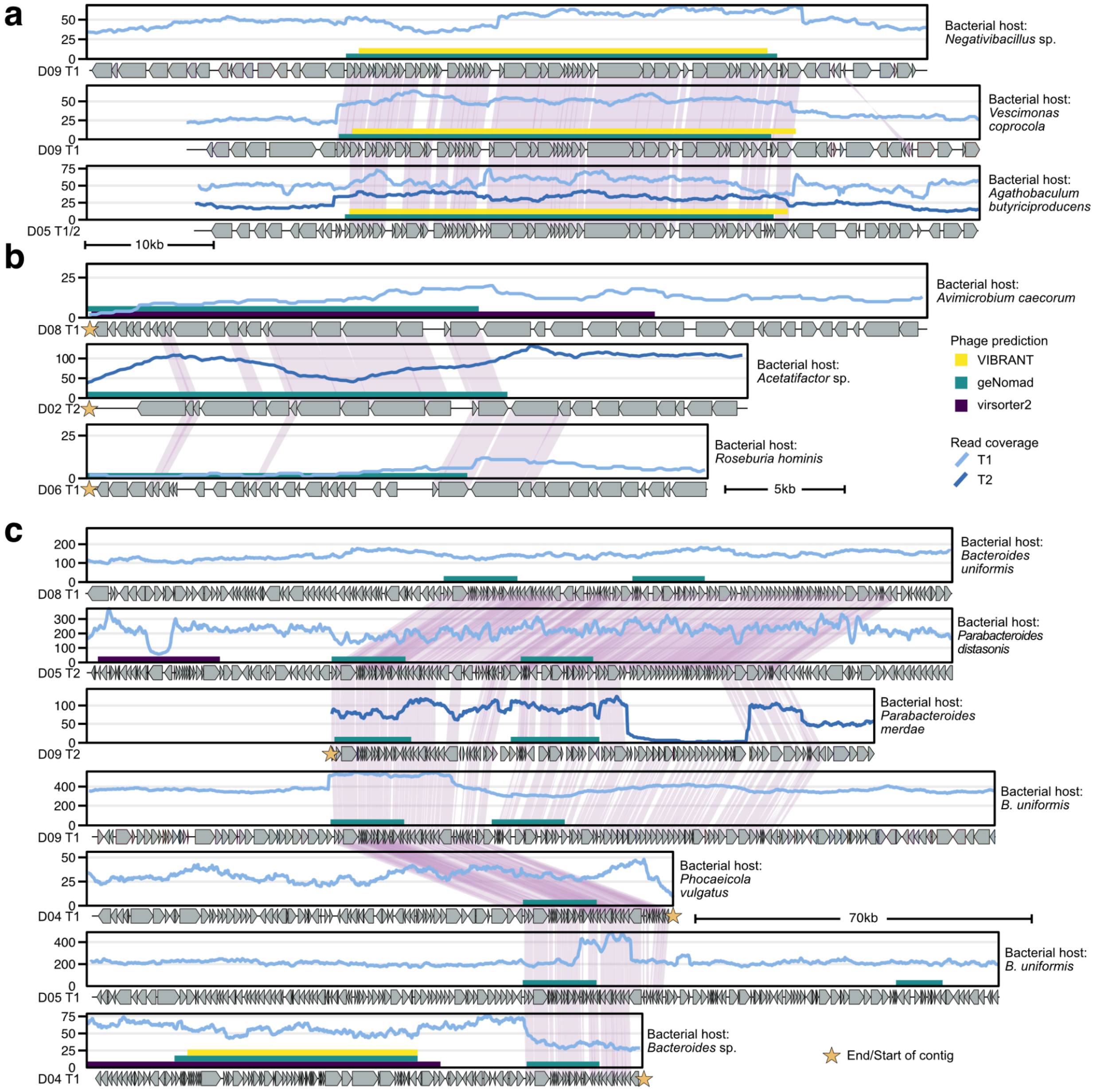


**Supplementary Figure 8: Coverage and synteny plots for clusters of phages assembled in taxonomically distinct host contexts.**

**a)** Coverage and synteny plots for a cluster of three phages that were assembled into three different host contexts. The host contexts were annotated as *Negativibacillus* sp. (in the family *Ruminococcaceae*), *V. coprocola* (in the family *Oscillospiraceae*), and *A. butyriciproducens* (in the family *Butyricicoccaceae*), all in the order *Oscillospirales*. **b)** Coverage and synteny plots for a cluster of three phages that were assembled into three different host contexts. The host contexts were annotated as *A. caecorum* (in the family *Ruminococcaceae* and the order *Oscillospirales*), *Acetatifactors* sp., and *R. hominis* (the latter two in the family *Lachnospiraceae* and the order *Lachnospirales*), all in the class *Clostridia*. **c)** Coverage and synteny plots for a cluster of seven phages that were assembled into multiple different host contexts. The host contexts were annotated as several *Bacteroides* species (in the family *Bacteroidaceae*), two *Parabacteroides* species (in the family *Tannerellaceae*), and *P. vulgatus* (in the family *Bacteroidaceae*), all in the order *Bacteroidales*. For all panels, the shaded regions between gene arrows indicate amino acid similarity greater than 80% and stars in the gene plots represent the start or end of assembled contigs.

Similarly, colored segments at the bottom of coverage plots indicate phage predictions from VIBRANT, geNomad, and virsorter2 across all panels.

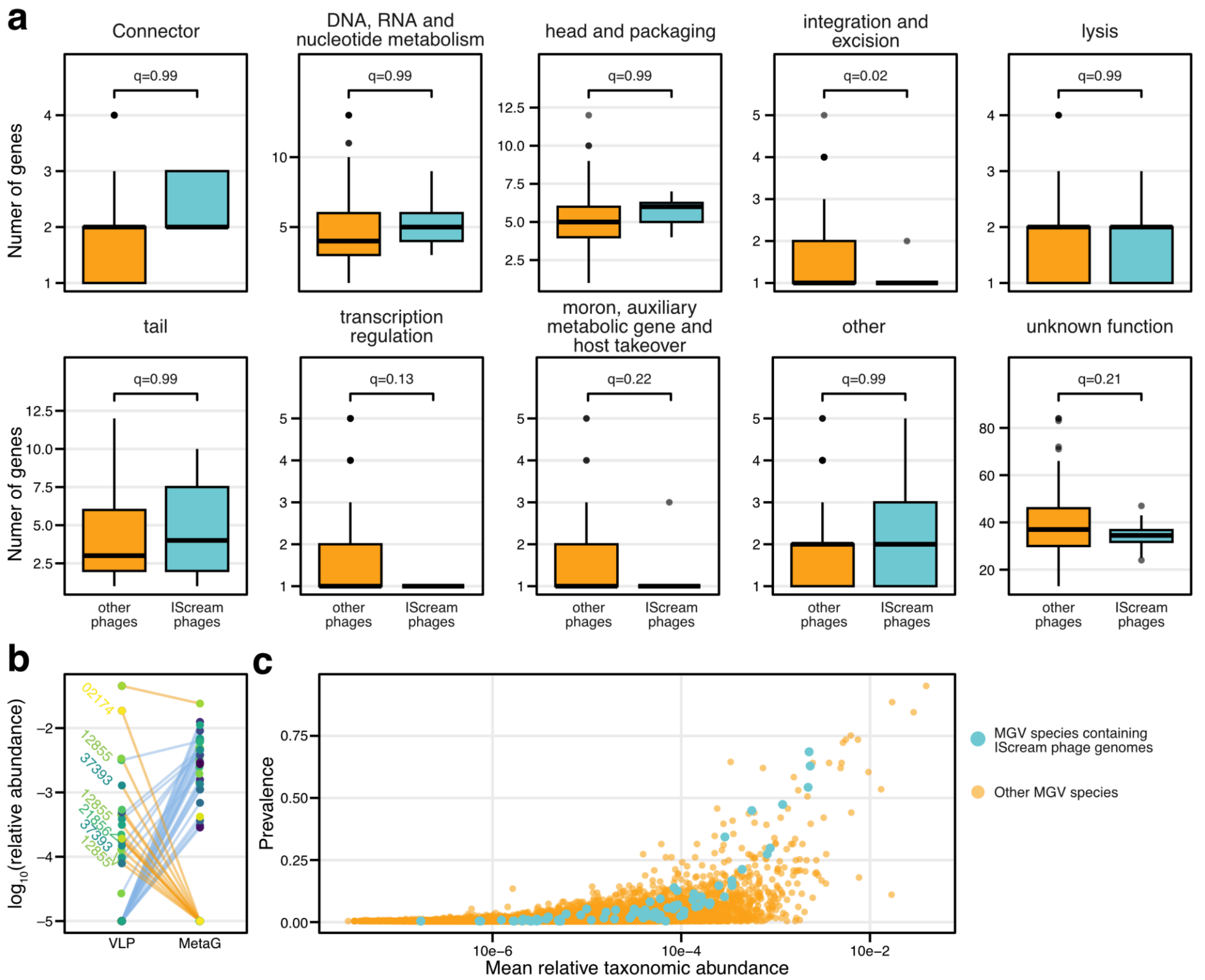


**Supplementary Figure 9: IScream phages have phage-like gene content and are present in MGV.**
**a)** Number of genes classified as different functional groups, identified from Pharokka, shown for both IScream phages and all other integrated phages. Both groups of phages were identified through structural variation analysis. Each panel is annotated by a Benjamini-Hochberg-corrected P-value, resulting from testing differences in gene numbers with a two-sided Wilcoxon test (Other phages n = 569, IScream phage n = 22) . Boxplots show the interquartile ranges (IQRs) as boxes, with the median as a black horizontal line, whiskers extending up to the most extreme points within 1.5-fold IQR, and outliers are indicated as dots. **b)** Log10-transformed relative taxonomic abundance for all phage species containing IScream phages identified in MGV genomes for data from Liang *et al.* (see **Methods**). Each dot represents a phage species within one individual, sequenced either in bulk metagenomic (MetaG) or virus-like particle (VLP)-enriched samples, and are connected by lines to indicate the same individual. Different phage species are indicated by different colors. **c)** Mean relative taxonomic abundance is plotted against prevalence for all phages identified by phanta in the metagenomes of healthy individuals from Yachida *et al.* (see **Methods**). Phage species containing IScream phages identified in MGV genomes are highlighted in cyan.

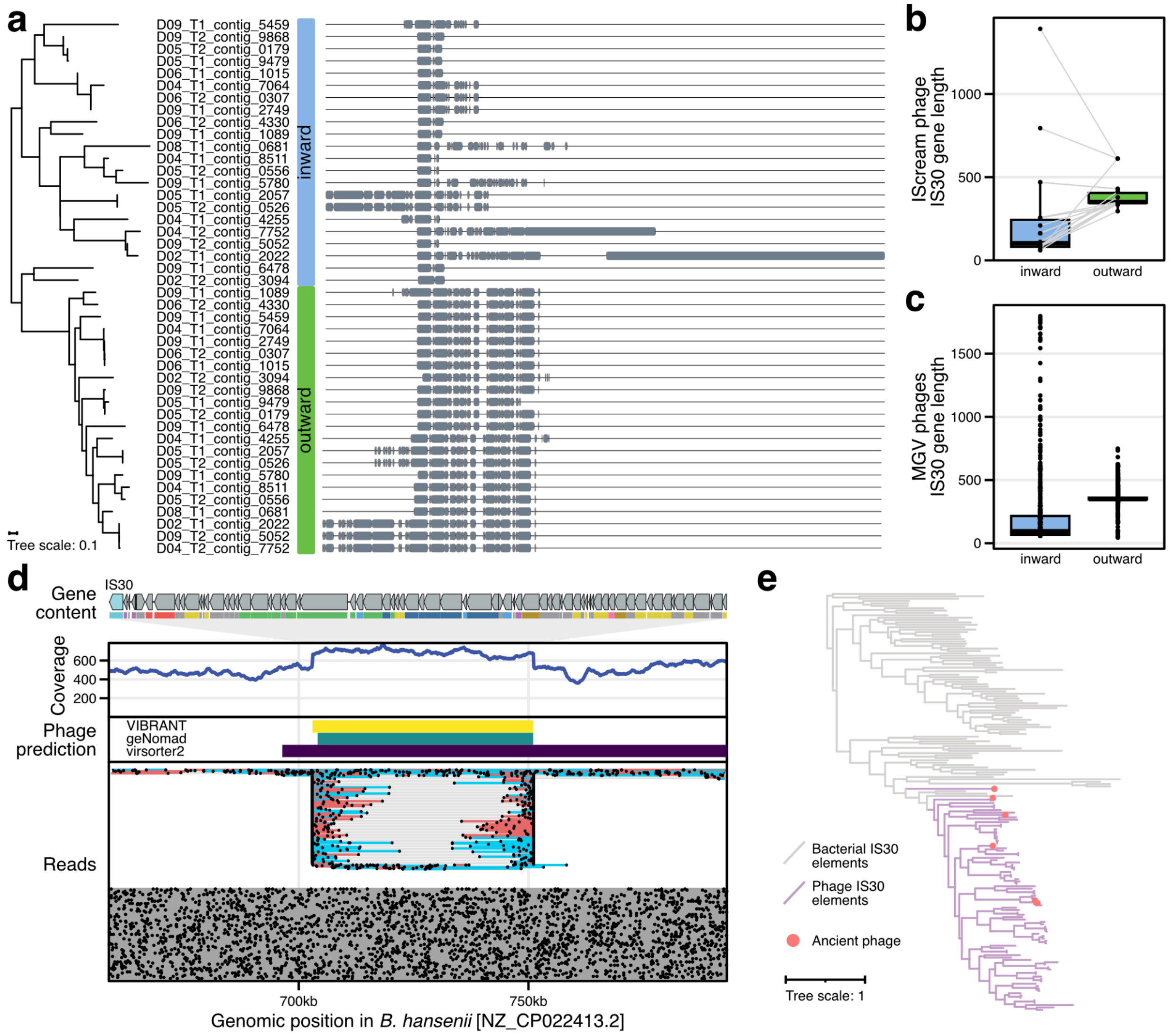


**Supplementary Figure 10: IS30 clustering and *Blautia hansenii* IScream phage.**
**a)** Multiple sequence alignment of all IS30 open reading frames in IScream phages with structural variation evidence, assembled in this study, visualized through the ete3 toolkit (boxes indicate alignments and empty areas indicate gaps in the alignment). The outward- and inward-directed IS30 proteins form two separate clusters, indicated by the tree reconstructed from the multiple sequence alignment. **b)** Boxplot showing the IS30 gene length for outward- and inward-directed IS30 genes from IScream phages assembled here. Boxplots show the interquartile ranges (IQRs) as boxes, with the median as a black horizontal line, whiskers extending up to the most extreme points within 1.5-fold IQR, and all data points are indicated as dots. **c)** Boxplot showing the IS30 gene length for outward- and inward-directed IS30 genes from all potential IScream phages identified in MGV. All boxplots show the interquartile ranges (IQRs) as boxes, with the median as a black horizontal line, whiskers extending up to the most extreme points within 1.5-fold IQR, and outliers indicated as dots **d)** Evidence for the presence of circular phage genomes from read alignments against the *B. hansenii* reference genome (NZ_CP022413.2). Read alignments are separated into linked alignments (top part) and single alignments (bottom part). Each line represents the alignment of a single read with dots showing the start and end of each alignment. Linked alignments are colored by strand and connected by a thin grey line to indicate that they originate from the same read. The remainder of this figure panel shows coordinates of phage predictions, read coverage, and the gene content of the phage genome (as determined by structural variation calling). **e)** Tree constructed from a multiple sequence alignment of IS30 proteins clustered at 70% amino acid identity (outward-directed IS30 proteins from MGV IScream phages, IScream phages assembled here, and bona-fide bacterial IS30 elements) together with IS30 proteins found overlapping phage predicted in assemblies of paleofeces.
